# The H2A.Z.1/PWWP2A/NuRD-associated protein HMG20A controls early head and heart developmental transcription programs

**DOI:** 10.1101/2022.05.04.490592

**Authors:** Andreas Herchenröther, Stefanie Gossen, Tobias Friedrich, Alexander Reim, Nadine Daus, Felix Diegmüller, Jörg Leers, Hakimeh Moghaddas Sani, Sarah Gerstner, Leah Schwarz, Inga Stellmacher, Laura Victoria Szymkowiak, Andrea Nist, Thorsten Stiewe, Tilman Borggrefe, Matthias Mann, Joel P. Mackay, Marek Bartkuhn, Annette Borchers, Jie Lan, Sandra B. Hake

## Abstract

Specialized chromatin-binding proteins are required for DNA-based processes during development. We recently established PWWP2A as direct histone variant H2A.Z interactor involved in mitosis and cranial-facial development. Here, we identify the H2A.Z/PWWP2A-associated protein HMG20A as part of several chromatin-modifying complexes including NuRD, and show that it localizes to genomic regulatory regions. Hmg20a depletion causes severe head and heart developmental defects in *Xenopus laevis.* Our data indicate that craniofacial malformations are caused by defects in neural crest cell (NCC) migration and cartilage formation. These developmental defects are pheno-copied in HMG20A-depleted mESCs, which show inefficient differentiation into NCCs and cardiomyocytes (CMs). Accordingly, loss of HMG20A caused striking deregulation of transcription programs involved in epithelial-mesenchymal transition (EMT) and cardiac differentiation, thereby providing insights into the regulatory circuits controlled by HMG20A. Collectively, our findings implicate HMG20A as part of the H2A.Z/PWWP2A/NuRD-axis and reveal it as a key modulator of the intricate developmental transcription programs that guide NCC and cardiomyocyte differentiation.

## Introduction

Proper control of chromatin structure is important for the regulation of eukaryotic gene expression, which is required for successful embryonic development and efficient stem cell differentiation. This complex process includes the incorporation of histone variants into chromatin, ATP-dependent remodelling of nucleosomes, the action of regulatory RNAs as well as chemical modifications of DNA and histone proteins. How histone variants, as important factors controlling the accessibility of the underlying genetic information^1^, mechanistically coordinate gene regulation is still not fully understood.

The histone variant H2A.Z is highly conserved^2^ and in vertebrates encoded by two genes (*H2AFZ* and *H2AFV*), whose protein products (H2A.Z.1 and H2A.Z.2.1) differ in only three amino acids^3^. In primates, the *H2AFV* RNA can be alternatively spliced giving rise to an additional H2A.Z.2.2 protein with a shortened C-terminus that destabilizes nucleosomes^4, 5^. Importantly, H2A.Z has been demonstrated to be essential for embryogenesis and is involved in neurodevelopmental processes^2, 6–8^. However, the underlying molecular mechanism(s) remains ill-defined, although H2A.Z has been implicated in many DNA-based processes including transcriptional regulation, cell cycle control and DNA repair^8–13^.

Together with recent studies, our previous work has shed some light on the H2A.Z functional network in gene regulation and embryonic development, where the interactomes of H2A.Z isoforms have been determined. We discovered, among several other proteins, PWWP2A as a direct and highly specific binder of H2A.Z.1 and H2A.Z.2.1 nucleosomes^14, 15^. Depletion of PWWP2A causes severe phenotypes, such as significant delays in mitotic progression in human cell lines and strong craniofacial defects in *Xenopus laevis*, most likely arising from defects in neural crest cell (NCC) migration and differentiation^14, 16^. We found that PWWP2A regulates many H2A.Z- controlled genes, probably by recruitment of chromatin-modifying proteins, such as an MTA1-specific core nucleosome and remodelling and deacetylase (NuRD) complex (M1HR). M1HR lacks the remodelling CHD sub-unit usually found in NuRD, due to competition between PWWP2A and the MBD proteins that recruit CHD to the NuRD complex^15, 17–19^.

Besides M1HR, PHD Finger protein 14 (PHF14), Retinoic Acid Induced 1 (RAI1), Transcription Factor 20 (TCF20) and High Mobility Box 20A (HMG20A) proteins were repeatedly identified in both H2A.Z.1/2.1 and PWWP2A interactomes^14, 15, 20^. A complex, which we termed PRTH due to the initials of the four complex members. These four proteins have previously been demonstrated to be repelled by histone H3 lysine 4 trimethylation (H3K4me3) and to be part of one unified complex^21^. More importantly, all four proteins have been shown to be deregulated or mutated in neurodevelopmental diseases including intellectual disabilities and autism spectrum disorders^22^ by disrupting the H3K4 methylation signature that ensures normal brain development^23^. As PWWP2A depletion in *Xenopus laevis* results in severe defects in craniofacial development, we wondered whether its association with PHF14/RAI1/TCF20/HMG20A might be – at least partially – causative for this observation. Hence, we turned our attention to the functional characterization of one member of this complex. In this study, we focused on HMG20A (previously termed iBRAF), as little is known about its histone variant-related function(s), particularly in embryonic development.

Here, we expand the published HMG20A’s interactome by taking advantage of GFP- HMG20A expressing human cells and identify, besides BHC/CoREST and PRTH proteins also NuRD complex members. Subsequently, we demonstrate that binding to those proteins depends on HMG20A’s C-terminal region, which contains a coiled-coil (CC) domain, while the N-terminus with the conserved HMG box conveys DNA-binding activity. Further, ChIP-seq experiments reveal a strong enrichment of HMG20A at two separate genomic regions: nucleosome-depleted transcriptional start sites (TSSs) surrounded by H2A.Z/PWWP2A-containing nucleosomes, and H2A.Z/PWWP2A- lacking intronic enhancer regions. Due to a high sequence conservation of HMG20A among human, mouse and *Xenopus*, we next determined the biological significance of HMG20A using *Xenopus laevis* wherein depletion of Hmg20a *in vivo* resulted in severe defects in craniofacial and heart development. In addition, we observed that both NCC migration and cartilage differentiation transcription processes were impaired, phenotypes that were also observed upon loss of Pwwp2a. Similar defects were recapitulated by Hmg20a-depleted mouse embryonic stem cells (mESCs), which showed compromised differentiation to neural crest cells (NCCs) and beating cardiomyocytes (CMs).

On the molecular level, we utilized CUT&RUN to genome-wide profile HMG20A chromatin binding in primed mESCs – a common point shortly before differentiation to NCCs or CMs. By this approach, we revealed the enrichment of HMG20A at regulatory regions that contain binding motifs highly similar to our findings in human cells. Additional gene expression analysis of Hmg20a-depleted NCCs and CMs revealed a striking deregulation of transcription programs including genes responsible for epithelial-mesenchymal transition (EMT) and cardiac differentiation. These findings suggest a mechanistic basis for the observed defects in head and heart development. In summary, our study identifies HMG20A as a novel chromatin binding protein that acts in conjunction with either within H2A.Z/PWWP2A/M1HR or within BHC/NuRD complexes that reside at promoter and enhancer regions. Through this biochemical activity, HMG20A serves as a key modulator participating in regulating important differentiation and developmental programs.

## Results

### HMG20A interacts with PRTH, BHC/CoREST and NuRD complexes

Previously, we employed label-free quantitative mass spectrometry (lf-qMS) approaches to identify H2A.Z- and PWWP2A-mononucleosome binding proteins in HeLa Kyoto (HeLaK) cell lines^14, 16, 20^. Among several other proteins, we detected the PRTH complex members PHF14, RAI1, TCF20 and HMG20A as strong binders in both H2A.Z and PWWP2A interactome data sets. To gain further insights into any functional H2A.Z connection between these proteins, we focused our attention on HMG20A, which is vertebrate-specific and contains an internal HMG box and a C-terminal coiled-coil (CC) region (Supplemental Figure 1A). First, we confirmed the association of H2A.Z with HMG20A by performing mononucleosome immunoprecipitations (mononuc-IPs) with HeLaK cells stably expressing GFP-H2A and GFP-H2A.Z.1 (Figure 1A). Next, we generated HeLaK cell lines stably expressing N-terminally GFP-tagged HMG20A (Supplemental Figure 1B, C). As expected, GFP-HMG20A was predominantly observed in the nucleus, with a localization pattern similar to that of endogenous HMG20A protein (Figure 1B).

**Figure 1:**
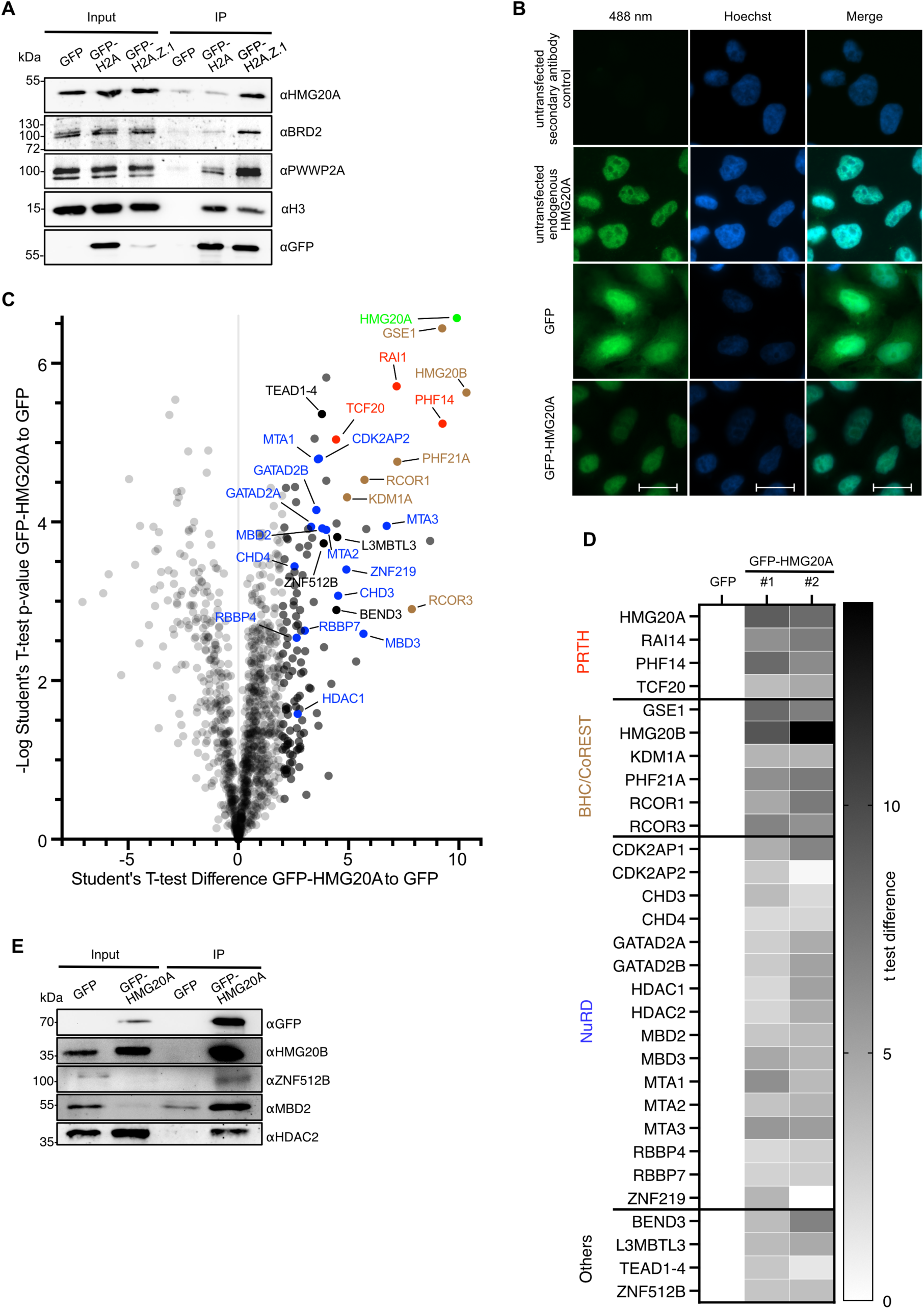
HMG20A binds repressive chromatin-modifying complexes. **A)** Immunoblots of GFP, HMG20A, BRD2 and PWWP2A (positive controls) as well as H3 upon GFP, GFP-H2A and GFP-H2A.Z.1 mono-nuc IPs. **B)** Immunofluorescence microscopy images of GFP, GFP-HMG20A and endogenous HMG20A proteins (488 nm, green) in HeLaK cells. DNA is stained with Hoechst (blue). Scale bar: 20 µm. **C)** Volcano plot of label-free interaction partners of GFP-HMG20A-associated mononucleosomes. Significantly enriched proteins over GFP-associated mononucleosomes are shown in the upper right part. t-Test differences were obtained by two-sample t-test. HMG20A is highlighted in bright green, PRTH members in red, BHC/CoREST members in brown, NuRD members in blue, other proteins in black and background binding proteins in grey. See also Supplemental Figure 1E for Volcano plot of second biological replicate and Supplemental Table 1 for detailed list of HMG20A binders. **D)** Heatmap of significant outliers from two independent GFP-HMG20A mononucleosome pull-downs analysed by lf-qMS (see C and Supplemental Figure 1E) normalized to GFP. Scale bar: log2-fold t-test differences. **E)** Immunoblots of GFP and GFP-HMG20A mononuc-IPs detecting endogenous members of the BHC/CoREST (HMG20B) and NuRD (MBD2, HDAC2) complexes as well as ZNF512B protein.

To identify the HMG20A interactome, we generated mononucleosomes (Supplemental Figure 1D) and immunoprecipitated those with GFP-TRAP beads; GFP- or GFP- HMG20A bound proteins were then subjected to on-bead tryptic digestion and then quantified by lf-qMS. Identification of the PRTH members PHF14, RAI1, TCF20 and the BHC/CoREST proteins HMG20B, GSE1, PHF21A, KDM1A, RCOR1 and RCOR3 as documented HMG20A binders^23, 24^ provided confidence in our approach (Figure 1C, D, Supplemental Figure 1E, Supplemental Table 1). The interaction of HMG20A with BHC/CoREST and NuRD members was independently verified by immunoblotting (Figure 1E). Interestingly, we detected several proteins and subunits of complexes previously identified to interact with both H2A.Z and PWWP2A but not uniquely with H2A.Z nucleosomes (Supplemental Figure 1F), implying that HMG20A might rather be a PWWP2A-associated protein rather than a direct binder of H2A.Z. Of particular interest was the observation that the complete NuRD complex was reproducibly pulled-down by GFP-HMG20A bound nucleosomes, as were members of the TEAD transcription factor family, the zinc finger protein ZNF512B and the chromatin modifying factors BEND3 and L3MBTL3 (Figure 1C, D, E, Supplemental Figure 1E, F). In brief, we identified PRTH, BHC/CoREST, and NuRD complexes and as well as several chromatin-modifying proteins as HMG20A binding factors. Some of these proteins are also part of the H2A.Z/PWWP2A interactomes (PRTH, M1HR, ZNF512B), while others appear to be HMG20A-specific (BHC/CoREST, NuRD), providing at least two alternative mechanisms for HMG20A to exert its biological function.

### HMG20A’s coiled-coil domain containing C-terminus is sufficient for NuRD binding

Having discovered that HMG20A interacts with the complete NuRD complex, we next asked how HMG20A contacts this complex. To answer this question, we co-expressed GFP-HMG20A with combinations of FLAG-tagged MTA1, MTA2, RBBP4 and/or HDAC1 in HEK293 cells. Immunoprecipitation of GFP-HMG20A from cell extracts revealed strong association with MTA1 but not MTA2 (Figure 2A). Additionally, we observed strong binding to HDAC1, while HMG20A did not interact with RBBP4 alone. Similarly, we detected interactions of GFP-HMG20A with HA-MBD2/3, -GATAD2A and FLAG-CHD4 (Figure 2B, top), indicating that HMG20A, in contrast to PWWP2A^17, 19, 25^, pulls down both remodelling and deacetylase subunits of NuRD. Further mapping of the interaction with CHD4 using deletion constructs (Figure 2B, bottom) revealed that the interaction with HMG20A is mediated by the central DNA translocase domain, rather than with the N- and C-terminal domains that are known to mediate interactions with nucleosomes, poly-ADP-ribose and GATAD2A/B^26, 27^ (Figure 2B, top).

**Figure 2:**
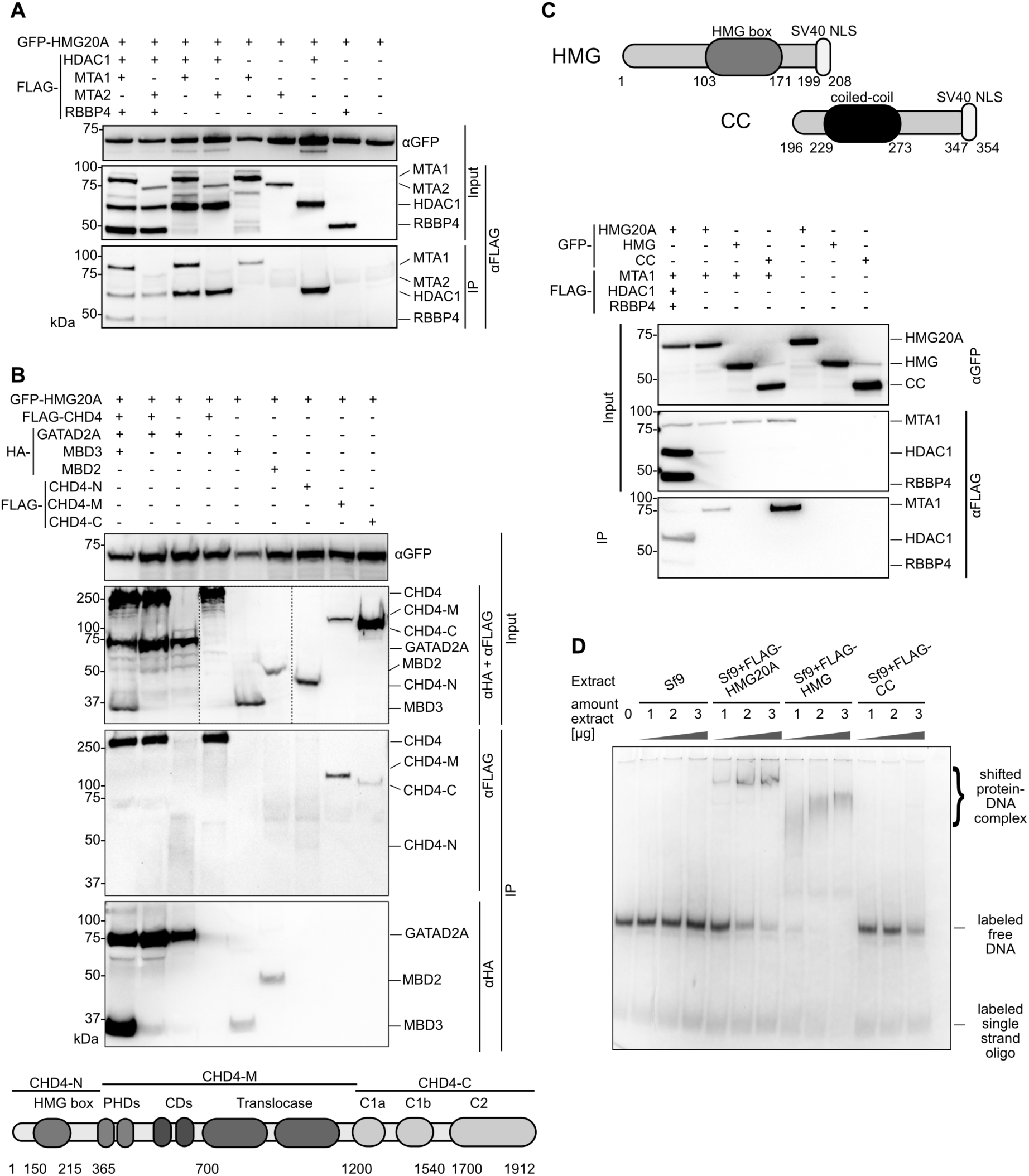
HMG20A binds NuRD complex components and DNA. **A)** Anti-GFP immunoprecipitations of HEK293 cell extracts co-transfected with GFP- HMG20A and FLAG-HDAC1, -MTA1, -MTA2 and -RBBP4. Proteins were detected with anti-FLAG or anti-GFP antibodies. **B)** Top: Anti-GFP immunoprecipitations of HEK293 cell extracts co-transfected with GFP-HMG20A and FLAG-CHD4 (CHD4), -CHD4-N-terminus (CHD4-N), -CHD4- middle domain (CHD4-M), -CHD4-C-terminus (CHD4-C) and HA-GATAD2A, -MBD2 and -MBD3. Proteins were detected with anti-FLAG and anti-HA or anti-GFP antibodies. Bottom: schematic depiction of CHD4 deletion constructs. **C)** Top: schematic depiction of HMG20A deletion constructs. Bottom: anti-GFP IPs of HEK293 cell extracts co-transfected with GFP-HMG20A and its deletions (HMG, CC) and of NuRD members (FLAG-MTA1, HDAC1-FLAG, FLAG-RBBP4). Proteins were detected with anti-FLAG or anti-GFP antibodies. **D)** EMSA of increasing amounts of extracts from Sf9 cells expressing FLAG-HMG20A and its deletions (see C, top) using Cy5-labelled DNA.

Next, we asked which region in HMG20A is required for NuRD binding. Like previously reported for the BHC/CoREST complex^28^, it is HMG20A’s C-terminal region containing the CC domain but not the HMG box-containing N-terminus (Figure 2C, top) that is required for its binding to MTA1 when expressed together with combinations of FLAG-tagged MTA1, RBBP4 and HDAC1 in HEK293 cells (Figure 2C, bottom).

Lastly, we wondered whether the HMG box from HMG20A retains its ability to bind DNA, given that it harbours the conserved amino acids known to be required for nucleic acid recognition by HMG domains^29^ (Supplemental Figure 2A). To test this possibility, we expressed FLAG-HMG20A and the corresponding deletion constructs (Figure 2C, top) in Sf9 cells (Supplemental Figure 2B and C) and performed Electromobility Shift Assays (EMSAs) using a Cy5-labelled, 100-bp random DNA probe. As expected, HMG20A’s N-terminal domain containing the HMG box is indeed capable of binding free DNA (Figure 2D), in agreement with a recent report where the full-length HMG20A protein was used^30^.

Together, these results reveal details of the interaction between HMG20A and NuRD components (CHD4 and MTA1) and further highlight the conserved functions of HMG20A’s N-terminal and C-terminal regions in DNA binding and protein-protein interaction, respectively. Our data lead to a working model by which HMG20A is involved in the recruitment of chromatin modifiers, such as NuRD, to specific genomic locations.

### HMG20A localizes to H2A.Z/PWWP2A-containing regulatory regions and H2A.Z- lacking intronic enhancer regions

Having revealed HMG20A’s DNA binding ability, we next mapped its genomic location by chromatin immunoprecipitation followed by high-throughput sequencing (ChIP-seq) (Supplemental Figure 3A), using HeLaK cells stably expressing GFP-HMG20A. We compared HMG20A sites with published ChIP-seq data for H2A.Z and PWWP2A^14, 16^ and found two clusters: (i) HMG20A binding sites that overlapped strongly with H2A.Z/PWWP2A occupancy and (ii) HMG20A binding sites that overlapped weakly with H2A.Z/PWWP2A (HMG20A-only). (Figure 3A top, Supplemental Figure 3A). Of the approximately 12,000 HMG20A sites, around 70% overlapped with H2A.Z and/or PWWP2A regions (HMG20A/H2A.Z/PWWP2A sites), whereas around 30% displayed reduced H2A.Z and PWWP2A presence (HMG20A-only sites) (Figure 3A bottom).

**Figure 3:**
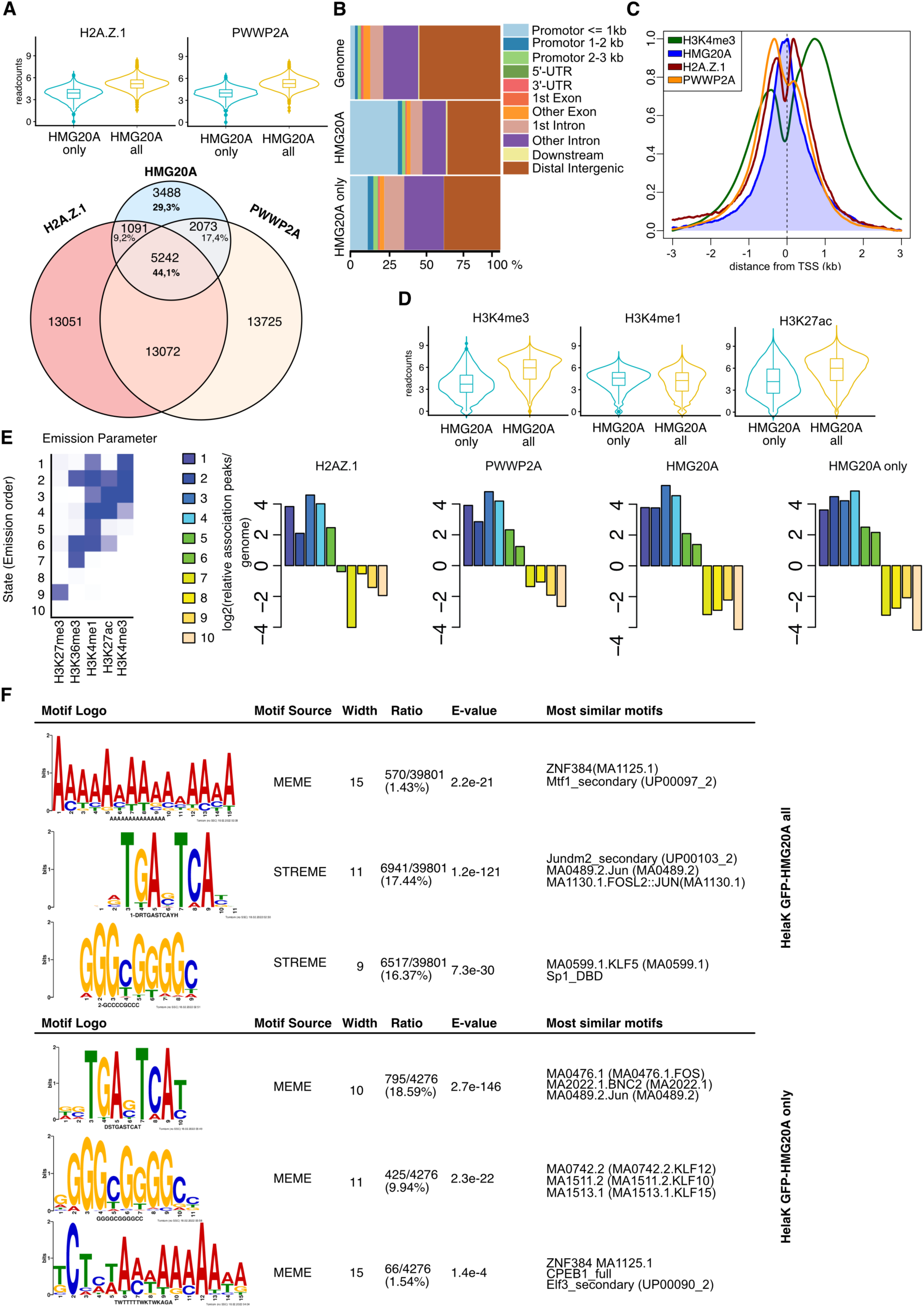
HMG20A localizes to distinct regulatory chromatin regions. **(A)** Top: Violin plots of GFP-H2A.Z.1 (left) and GFP-PWWP2A (right) ChIP-seq read counts at GFP-HMG20A binding sites. Bottom: Venn-Diagram displaying numbers of GFP-H2A.Z.1, -PWWP2A and -HMG20A ChIP-seq binding sites and their overlap. **(B)** Enrichment plot representing genomic regions after GFP-HMG20A ChIP-seq. **(C)** Average binding profiles across transcriptional start sites of GFP-HMG20A (blue), -H2A.Z.1 (red), -PWWP2A (orange) and H3K4me3 (green) mean coverage signals at TSS of expressed genes. **(D)** Violin plots of H3K4me3 (left), H3K4me1 (middle) and H3K27ac (right) ChIP-seq read counts at GFP-HMG20A binding sites **(E)** Left: ChromHMM^32, 33^-based characterization of chromatin states of GFP-HMG20A- containing genomic regions. Heatmap depicts emission parameters of the HMM and describes the combinatorial occurrence of individual histone modifications in different chromatin states. Right: Chromatin-state enrichment of HMG20A-, PWWP2A-, H2A.Z.1- and HMG20A-only-enriched sites calculated to frequency in complete genome. **(F)** Top: Top-enriched motifs of all HMG20A ChIP-seq peaks identified with MEME and STREME analysis. Bottom: Top-enriched motifs in HMG20A-only ChIP-seq peaks identified with MEME analysis.

A subset of ChIP-seq enrichment sites were further validated by ChIP-qPCR (Supplemental Figure 3B). Intriguingly, the validated targets showed a stronger association of HMG20A with HMG20A-only sites compared with H2A.Z/PWWP2A- containing ones. Such a difference might be due to the fast turnover rate of H2A.Z that is typically observed in promoter regions^31^. Indeed, in the presence of H2A.Z/PWWP2A, HMG20A was found to be particularly enriched at regulatory regions, such as promoters, while HMG20A was favourably enriched at introns in the absence of H2A.Z/PWWP2A (Figure 3B). A closer examination of HMG20A localization at promoters revealed enriched binding of HMG20A to nucleosome-depleted regions (NDRs) at transcriptional start sites (TSSs) that are surrounded by PWWP2A-bound H2A.Z-containing +1 and -1 nucleosomes (Figure 3C). It is also notable that for all HMG20A binding sites (regardless of the presence or absence of H2A.Z/PWWP2A), it was only full-length HMG20A, but neither the N-terminal nor the C-terminal region alone, that was able to pull down chromatin (Supplemental Figure 3B), indicating that both DNA and HMG20A-interacting proteins are required for efficient chromatin binding (see Figure 2C, D).

To further characterize and distinguish HMG20A/H2A.Z/PWWP2A and HMG20A-only sites across the genome, we used available ENCODE data sets to define the presence of H3K4me3 as promoter, H3K4me1 as enhancer and H3K27ac as active regulatory marks. As depicted in Figure 3D, HMG20A-only sites are biasedly less marked by H3K4me3 and H3K27ac, while being mildly more H3K4me1-associated, supporting that idea that HMG20A-only sites are rather located in enhancers than promotors. To increase our confidence and better characterize genomic HMG20A binding regions, we performed a more powerful comparison between our datasets and chromatin states defined by training a 10-state model on ENCODE using ChromHMM^32, 33^ (Figure 3E, left). As shown in Figure 3E (right), HMG20A/H2A.Z/PWWP2A sites showed indeed a positive correlation with H3K4me3 and a negative correlation with the repressive H3K27me3 mark. Interestingly, HMG20A-only sites were additionally enriched with H3K36me3 (transcribed gene bodies), suggesting that HMG20A is associated with two main chromatin contexts: i) HMG20A together with H2A.Z and PWWP2A at promoters enriched with H3K4me3 and H3K27ac and ii) HMG20A-only sites at H3K4me1- enriched intronic enhancers within H3K36me3-positive genes.

Next, we used STREME and MEME^34^ to ask whether the HMG20A enriched sites contain a specific consensus sequence. As shown in Figure 3F, we found an overrepresented A-rich motif, which resembles the published HMG-box binding motif AGAACAAGAAA^35^. Interestingly, additional representative sequences such as the promoter-associated Fos/Jun, SP1 and KLF5 motifs were also detected. Given that Fos/Jun, SP1 and Klf5 were recently identified as potential regulators controlling the balance between mesendoderm and ectoderm during embryonic stem cell (ESC) differentiation^36^, this finding indicates a potential function of HMG20A in developmental processes. Accordingly, RNA-seq of HeLaK cells upon HMG20A knock-down with a specific siRNA pool determined only few genes to be affected in their transcription profile, with just 81 up- and 77 down-regulated genes (Supplemental Figure 3C and D, Supplemental Table 2).

Taken together, our data show HMG20A to localize to the NDRs within TSSs of H2A.Z- and PWWP2A-surrounded promoter nucleosomes as well as intronic enhancer regions within transcribed genes containing defined consensus sequences.

### HMG20A is required for craniofacial and heart development in *Xenopus laevis*

Our data so far hinted at HMG20A playing a role in developmental processes. Hence, we turned to the African clawed frog *Xenopus laevis* as an animal model, which is easily manipulated. The Xenopus Hmg20a homologue displays high conservation in both the HMG box and the coiled-coiled region (Supplemental Figure 4A). Whole-mount RNA *in situ* hybridization of various developmental stages as well as RT-qPCR analyses detected endogenous *X. laevis hmg20a* mRNA at all time points (Supplemental Figure 4B). Maternal expression of *hmg20a* is seen at early cleavage and blastula stages (Supplemental Figure 4C-F); zygotic expression is high at gastrula stages, where *hmg20a* is ubiquitously expressed (Supplemental Figure 4G). At neurula stages, *hmg20a* is detected in neural folds and cranial NCCs (Supplemental Figure 4H-J), followed by expression in migratory cranial NCCs, brain and eyes at tadpole stages (Supplemental Figure 4K-R).

To investigate the developmental role of *X. laevis* Hmg20a we performed loss-of-function analyses. A translation blocking antisense Morpholino oligonucleotide (MO) targeted against the *hmg20a* RNA (*hmg20a MO*) or a control MO (*co MO*) was injected, in combination with *lacZ* RNA as a lineage tracer into one blastomere at the two-cell stage. At tadpole stages, morphants showed craniofacial and eye defects as well as hyperpigmentation on the injected side (Figure 4A, B), for the most part indicative of defects in NCC development. To further analyse NCC migration, we performed *in situ* hybridization of *hmg20a MO-*injected embryos and controls using the NCC marker *twist*. Indeed, we observed a reduction of twist-positive migratory NCCs on the *hmg20a MO-*injected side (Figure 4C, D), which could be partially rescued by co-injection of human *HMG20A* cDNA (Figure 4C, D). As NCCs contribute to cranial cartilage formation, we performed collagen II immunostaining to visualize the cartilage of morphant and control embryos. Consistent with a function of Hmg20a in NCC migration, we observed a reduction of cartilage structures on the *hmg20a MO*-injected side compared to embryos injected with the *co MO* (Figure 4E, F). Again, these defects were partially rescued by co-injection of human *HMG20A* cDNA (Figure 4E, F). Interestingly, the expression pattern as well as the NCC-specific loss-of-function defects of *Xenopus hmg20a* are highly reminiscent of the ones previously observed for *Xenopus* Pwwp2a^14^, suggesting a partial functional overlap between these associated proteins.

**Figure 4:**
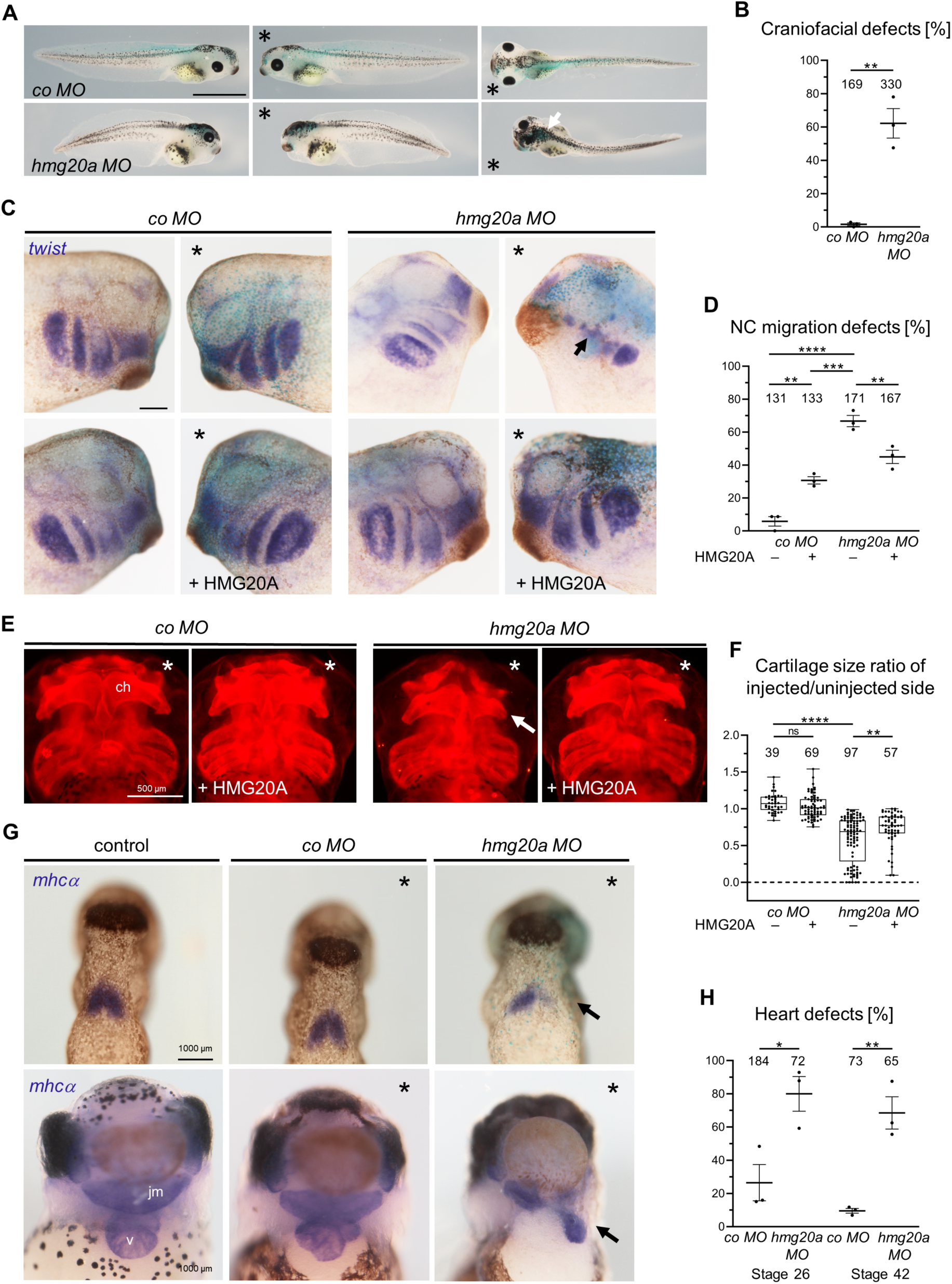
HMG20A depletion leads to craniofacial and heart malformations in frog. **(A)** Loss of function of Hmg20a leads to craniofacial and pigmentation defects in *Xenopus* tadpoles. Embryos were injected with 10 ng MO in combination with 80 pg *lacZ* RNA in one blastomere at the two-cell stage, * marks the injected side, white arrow marks pigmentation defects. Scale bar = 1 mm. **(B)** Graph summarizing the mean percentage of craniofacial defects in three independent experiments ± s.e.m. Number of embryos are indicated for each column. **p<0.01 (two-tailed unpaired Student’s t-test). **(C)** Hmg20 loss-of-function caused defects in cranial NC migration that can partially be rescued by co-injection of human HMG20A DNA. Embryos were injected with 10 ng MO in combination with 80 pg *lacZ* RNA (seen in blue) and analysed by *twist in situ* hybridization (seen in purple). For rescue experiments 130 pg HMG20A DNA was con-injected. * marks the injected side, arrow shows defect in cranial NC migration. Scale bar = 1mm. **(D)** Quantification of NC migration defects of three independent experiments injected and analysed as shown in (C). The data is presented as mean ± s.e.m., the number of embryos is indicated for each column. **p < 0.01, *** p < 0.001 **** p < 0.0001 (one- way ANOVA, Tukey’s multiple comparisons test). **(E)** Hmg20a-depleted *Xenopus* tadpole embryos show defects in cartilage formation (arrow). Embryos were injected with 10 ng MO in combination with 80 pg *membraneRFP (mbRFP)* RNA and analysed by collagen II immunostaining. For rescue experiments 100 pg HMG20A DNA was co-injected, * marks the injected side. Scale bar = 500 µm. **(F)** Box and whiskers plots summarize cartilage defects of at least three independent experiments analysed as in (E) and quantified by measuring the area of the ceratohyale cartilage. Number of embryos (n, above each bar) and median are indicated. The box extends from 25th to 75th percentile, with whiskers maximum at 1.5 IQR. **p < 0.01, **** p < 0.0001, ns.: not significant (one-way ANOVA, Tukey’s multiple comparisons test). **(G)** Hmg20a loss-of-function causes heart defects. Embryos were injected as in (C) and analysed by *mhcα in situ* hybridization. Top: Hmg20a-depleted tadpole stage 26 embryos show defects in the formation of the first heart field (arrow), while controls are unaffected. Bottom: At stage 42, Hmg20a depletion disturbs the three-chambered heart structure consisting of two atria (a) and a ventricle (v); the malformed heart is displaced towards the injected side. The jaw muscle (jm), which is also marked by *mhcα,* is also reduced in Hmg20a-depleted embryos. Scale bar = 1 mm. **(H)** Graph summarizing three independent experiments as shown in (G), data is presented as mean ± s.e.m., number of embryos is indicated for each column. *p < 0.05 (two-tailed unpaired Student’s t-test).

*Hmg20a*-depleted tadpoles also showed defects in heart morphology (Figure 4G, H), prompting us to analyse its role in heart development. By repeating the MO knockdown experiment, we traced expression of the cardiac differentiation marker *myosin heavy chain alpha* (*mhcα*) through *in situ* RNA hybridization. As shown in the top panel of Figure 4G, *hmg20a MO* injected embryos showed a reduction of *mhcα* in the first heart field compared to controls in earlier tadpole stages. Later at stage 42, control and *co MO*-injected embryos showed the typical heart structure consisting of two atria and one ventricle (Figure 4G bottom, H). However, this three-chambered heart structure could not be distinguished in *hmg20a MO* injected embryos. Furthermore, the malformed hearts were displaced toward the side of injection.

Taken together our data suggest that *Xenopus* Hmg20a plays an essential role in craniofacial and heart morphogenesis.

### Hmg20a drives NCC and cardiomyocyte differentiation of mESCs

Since Hmg20a depletion in *X. laevis* resulted in severe craniofacial and heart defects (see Figure 4), we speculated that HMG20A function is evolutionarily conserved and such defects would be recapitulated in the mammalian system. Hence, we first generated viable *Hmg20a* KO mESCs using a CRISPR/Cas9-based approach (Figure 5A, Supplemental Figure 5A, B). We differentiated^37^ WT and *Hmg20a* KO mESCs into NCCs (Figure 5B, Supplementary Figure 5C), mimicking the developmental process intimately related to the formation of craniofacial structures. Compared with WT cells, *Hmg20a* KO mESCs inefficiently differentiated to NCCs accompanied by a probable reduced migration ability, as assessed by RT-qPCR analysis of several NCC and/or EMT marker genes (Figure 5C).

**Figure 5:**
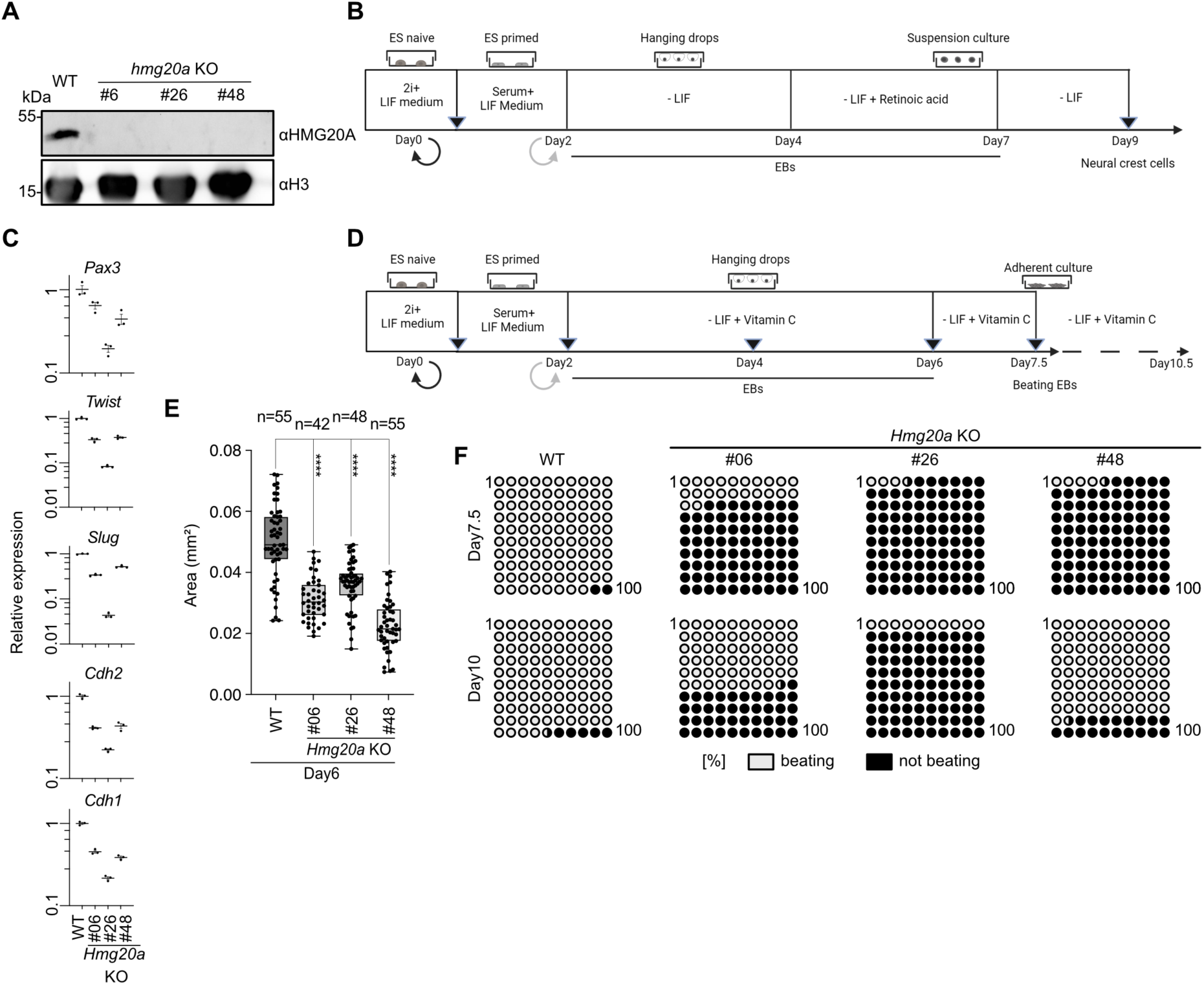
HMG20A is essential for cardiomyocyte and neural crest differentiation in mESCs. **(A)** HMG20A immunoblot of extracts from WT and three *Hmg20a* KO mESC clones. **(B)** mESC neural crest differentiation scheme. **(C)** RT-qPCR of neural crest and EMT marker genes in WT and 3 individual *Hmg20a* KO clones at Day9 of neural crest differentiation protocol normalized to *Hprt, 16S RNA and Gapdh* expression. **(D)** mESC cardiomyocyte differentiation scheme. **(E)** Size of embryoid bodies (EBs) of WT and three individual *Hmg20a* KO cells at Day6 of cardiomyocyte differentiation protocol. Number of measured EBs indicated above, **** p < 0.001, (two tailed Mann Whitney test). **(F)** Depiction of percent beating (gray) or non-beating (black) mESCs at Day7 (Top) or Day10 (bottom) of the cardiomyocyte differentiation procedure (see D).

Due to the observed reduction of *mhcα* expression in Hmg20a-depleted *X. laevis*, we next focused on the differentiation^38^ of WT and *Hmg20a* KO mESCs into beating CMs (Figure 5D, Supplementary Figure 5D). As a result, differentiation of mESCs to CMs was also compromised upon loss of Hmg20a, which was manifested by significantly smaller embryoid bodies (EBs) at the intermediate stage (Figure 5E, Supplemental Figure 5E). More importantly, *Hmg20a* KO mESCs failed to timely generate beating CMs on differentiation Day7.5 as compared with WT cells (Figure 5F top). Instead, some more *Hmg20a* KO CMs started beating beginning from Day10 (Figure 5F bottom), suggesting that loss of HMG20A does not lead to a complete halt of cardiomyocyte differentiation but rather results in a severe delay. Of note, the heart formation in mice is restricted to a short time window, between its onset in the primary heart tube at E8.5, and the final emergence of a four-chambered heart at E10^39^. Therefore, it is reasonable to infer that such a delay *in vivo* could give rise to a structural defect during heart formation, which would link the compromised *in vitro* differentiation to the *in vivo* observed defect.

Hence, our *in vitro* mESC model pheno-copies the defects observed *in vivo* in *Xenopus*, corroborating our findings that Hmg20a is functionally conserved and required for proper NCC and beating CM differentiation.

### Hmg20a regulates genes involved in NCC and cardiomyocyte differentiation

To better understand the molecular cause of the observed phenotypes, we first examined the genome-wide localization of Hmg20a by performing CUT&RUN-seq in WT and *Hmg20a* KO mESCs at the primed stage, which resembles an important join point before the separate differentiation into NCCs or CMs. As shown in Figure 6A, we identified a total of 2,545 *bona fide* Hmg20a-binding peaks, corresponding to 2,094 genes (Supplementary Table 3). Peaks were generally characterized by the absence of specific signals in the KO cell line. Examples of Hmg20a peaks are shown in Supplementary Figure 6A. Hmg20a peaks were mostly located in promotor regions, similar to our observations in HelaK cells (Figure 6B, Figure 3B). Subsequent analyses again revealed a strong enrichment of Jun/Fos, Klf and AT-rich motifs (Figure 6C). Hence, both Hmg20a genomic distribution and its consensus motifs are conserved between human HeLaK and mESC cells (see Figure 3B, F, Figure 6B, C). Also, GO analysis of Hmg20a-binding targets strongly supports our notion that Hmg20a is intimately involved in developmental processes, probably by modulating defined transcription programs (Supplementary Figure 6B).

**Figure 6:**
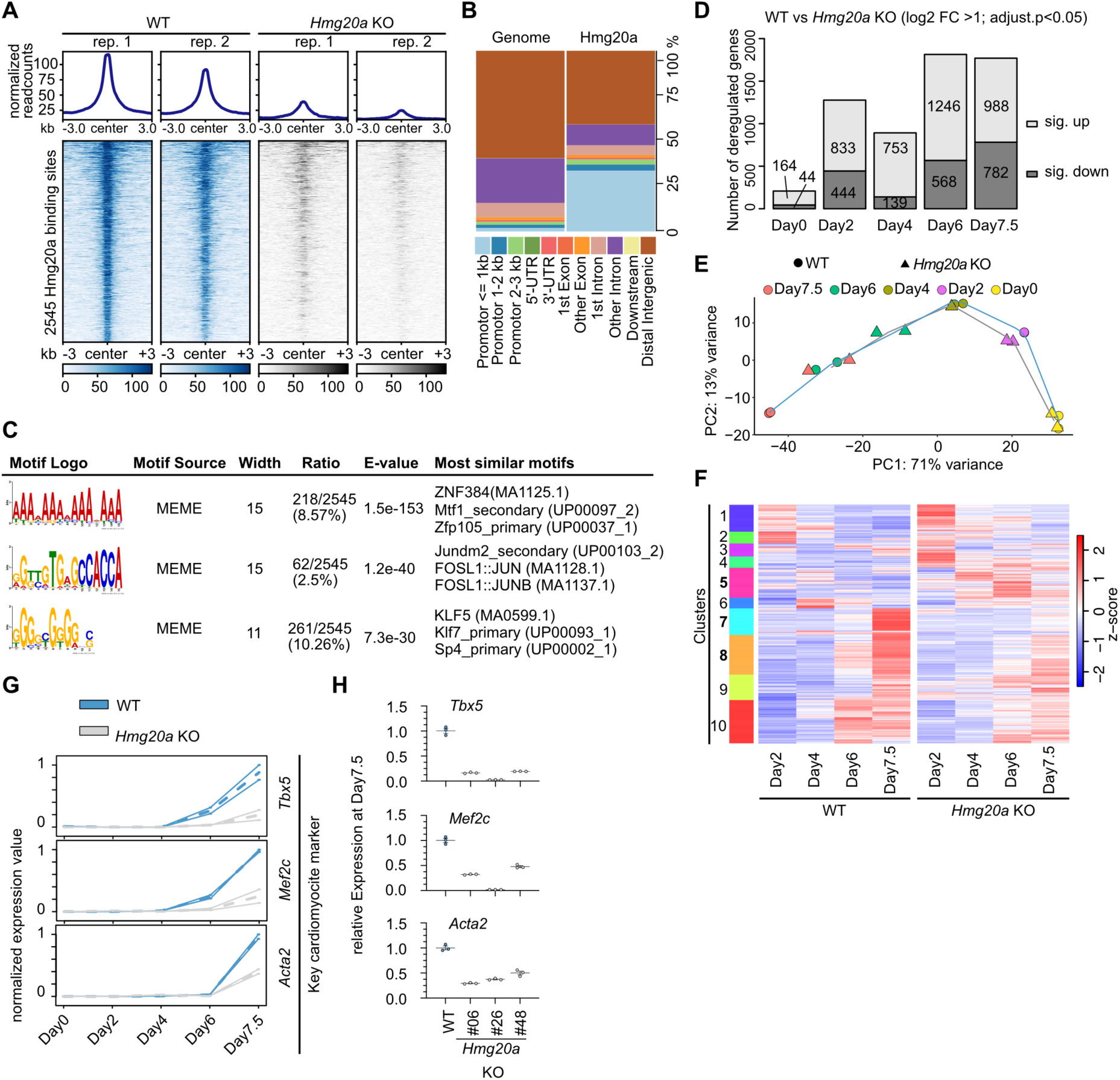
HMG20A regulates cardiomyocyte transcription programs. **(A)** Density heatmap of 2,545 Hmg20a binding sites detected in CUT&RUN. Color intensity represents normalized and globally scaled tag counts **(B)** Enrichment plot representing genomic regions of Hmg20a biding sites in CUT&RUN. **(C)** Top-enriched motif of Hmg20a CUT&RUN peaks identified with MEME and analysis. **(D)** Stacked Bar plot of numbers of significantly up (log2 fold change >1) and down (log2 fold change <-1) regulated genes (p<0.05) during indicated timepoints of cardiomyocyte differentiation **(E)** Principle component Analysis (PCA) of RNA-seq data of two replicates of WT (circle) and *hmg20a* KO clone#26 (triangle) at Day0 (yellow), Day2 (magenta), Day4 (olive), Day6 (green) and Day7.5 (red) differentiation time points (see Figure 5A for cardiomyocyte differentiation scheme). **(F)** Heat map showing the z-scaled expression values from all significant deregulated genes comparing the differentiation steps (Day 2 vs. 4; 4 vs. 6; 6 vs. 7.5). Genes are clustered according to the Euclidean distance by an unsupervised agglomerative hierarchical approach. Shown are the mean z-scales of 2 replicates for each day for wild type (left panel) or *Hmg20a* knock out (right panel) cells. **(G)** Line plots show the min-max normalized expression values for known cardiomyocyte marker genes at various differentiation time points. Plotted are mean expression values (dotted line) and the standard deviation (continuous line) for wild type (blue) or *Hmg20a* knock out (grey) cells. **(H)** RT-qPCR of cardiomyocyte marker genes in WT and three individual *Hmg20a* KO clones at Day7.5 of cardiomyocyte differentiation protocol normalized to *Hprt, 16S RNA and Gapdh* expression.

To examine this hypothesis, we first monitored Hmg20a-dependent transcriptome changes occurring at different time points during mESC to CM differentiation. RNA-seq experiments were performed on naïve (Day0), primed (Day2), two EB (Days4 and 6) and CM (Day7.5) stages of WT and *Hmg20a* KO clone #26 cells (see Figure 5D). Over time, an increasing number of genes deregulated in *Hmg20a* KO cells were detected (Figure 6D). Principal Component Analysis (PCA) revealed a striking stage-dependent gene expression trajectory (Figure 6E). *Hmg20a* KO Day7.5 cells showed a transcriptome profile comparable to that of Day6 WT cells, mirroring on the transcriptome level the severe developmental delay observed during differentiation of *Hmg20a* KO mESCs to CMs. To further analyse these differences, we examined changes of gene activity over time in WT, dividing them into 10 clusters according to their behaviour over time and compared them to the *Hmg20a* KO scenario (Figure 6F). Besides others, we identified 3 clusters with critical functions for CM differentiation that showed abnormal behaviour upon *Hmg20a* depletion (Figure 6F, Supplemental Figure 6C). Examples of key cardiomyocyte markers over time in CM differentiation are shown in Figure 6G. Although a slight clonal effect was observed between three *Hmg20a* KO clones, expression changes were validated by RT-qPCR for core cardiac TFs (Mef2c, Tbx5,)^40^ and the myofibroblast/cardiomyocyte marker Acta2^41^ (Figure 6H, Figure 5F), thereby reinforcing our observation of Hmg20a’s role in regulating CM transcription programs.

Together, these results indicate that Hmg20a is a key player in regulating lineage-specific differentiation transcription programmes.

## Discussion

Our work provides a deeper understanding of HMG20A function in development and transcriptional regulation of differentiation processes. The interactome of HMG20A revealed the known BHC/CoREST and PRTH complex members as well as all subunits of the NuRD complex and other chromatin-modifying proteins. We characterized the functional domains of HMG20A responsible for NuRD interaction and DNA binding and its occupancy at regulatory regions on chromatin. We uncover HMG20A’s conserved function in NCC-linked head and CM-linked heart development by directly regulating transcription and chromatin-regulatory factors driving lineage commitment or cell fate decision. We propose a working model where HMG20A acts together with either M1HR/PRTH at H2A.Z/PWWP2A surrounded promoters or with BHC/CoREST/NuRD complexes at intronic enhancers in transcribed genes (Figure 7, top). During development, HMG20A serves as a key modulator of faithful differentiation of stem cells into neural crest cells and cardiomyocytes by guiding lineage commitment programs (Figure 7, bottom).

**Figure 7:**
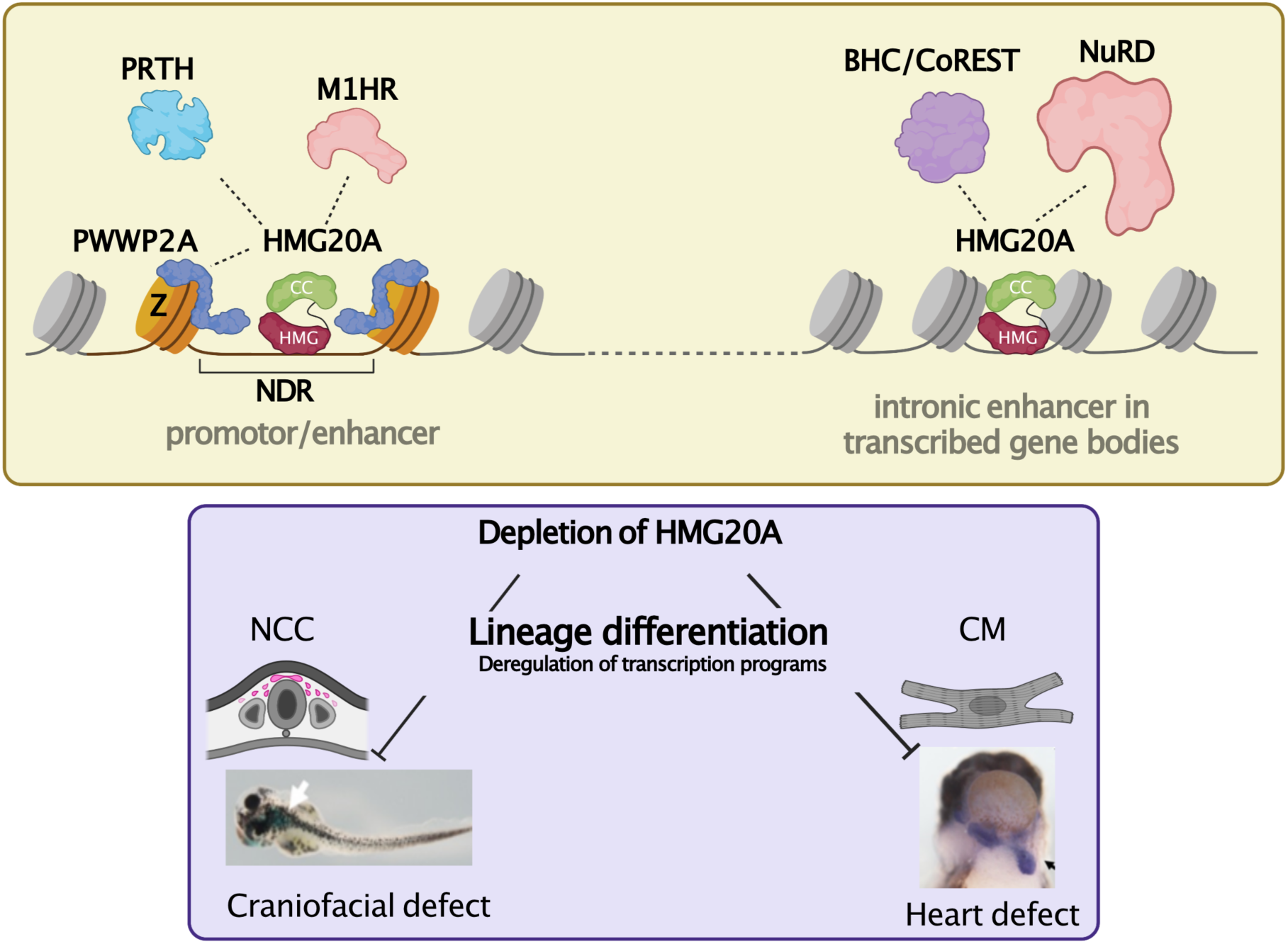
Model of HMG20A’s function in chromatin and transcriptional regulation during development. Top: HMG20A resides at two distinct regulatory regions: H2A.Z- and PWWP2A- surrounded nucleosome depleted regions (NDR) at promoter or intronic enhancer sites. Based on mass spectrometry data, we speculate that HMG20A binds to PRTH and M1HR components found at promoters/enhancers and BHC/CoREST and NuRD complexes at active intronic enhancers. Bottom: Depletion of Hmg20a in *Xenopus laevis* and mESCs leads to respective craniofacial and heart or neural crest cell (NCC) and cardiomyocyte (CM) differentiation defects, due to deregulated transcription programs.

### HMG20A is a member of several repressive chromatin-modifying complexes and binds to two distinct types of regulatory chromatin regions

The binding partners of HMG20A can be divided into two groups. The first group comprise BHC/CoREST, a known HMG20A associated complex identified by us and others^28^, the complete NuRD complex and several chromatin-modifying proteins. Members of this group are not part of the H2A.Z.1 and PWWP2A interactomes. The second group comprises proteins that are also associated with H2A.Z/PWWP2A, namely PRTH complex members and the MTA1, HDAC and RBBP core components (M1HR) of the NuRD complex. To fully understand the functional importance of HMG20A in lineage commitment it should be taken into consideration whether a particular HMG20A-interacting complex associates with H2A.Z.1/PWWP2A or not. Our ChIP-seq findings show that HMG20A binds to two distinct types of chromatin loci. On the one hand, HMG20A is correlated to H2A.Z/PWWP2A-cobound promoters. On the other hand, it is found at intronic enhancers in the absence of H2A.Z/PWWP2A. We observed that in contrast to PWWP2A, which exclusively interacts with the M1HR sub- complex of NuRD, the PWWP2A interactor HMG20A is surprisingly able to pull-down components of both deacetylase and remodelling subcomplexes (i.e. HDAC and CHD4) of NuRD. Recent ChIP-seq data in the rhabdomyosarcoma RH4 cell line similarly observed HDAC, MTA, RBBP subunits of NuRD at both active promoters and enhancers, but observed CHD only at enhancers^42^. Thus, it is tempting to speculate two working contexts for HMG20A, one of which associates with the M1HR complex at H2A.Z/PWWP2A-cobound promoters, and a second at intronic enhancer sites where it interacts with the complete NuRD complex, the BHC/CoREST complex and possibly also TEAD, L3MBTL3 and BEND3, as those are also not found in H2A.Z or PWWP2A interactomes (Figure 7).

Additionally, given the highly similar motifs found in both HMG20A-only and HMG20A/H2A.Z/PWWP2A binding sites (Figure 3F) and the failure of the HMG box-containing N-terminus to pull down chromatin (Supplemental Figure 3B), the DNA sequence alone is unlikely to be the primary factor determining the chromatin targeting of HMG20A. Hence, how the interacting proteins, such as BHC/CoREST, NuRD subunits, other PRTH members, H2A.Z, PWWP2A and others, assist in HMG20A targeting to distinct regulatory regions or not will require additional investigations.

### Hmg20a controls NCC and CM differentiation programs

We found Hmg20a to be required for proper craniofacial and heart formation during embryonic development. The observed defect in craniofacial formation is likely linked to early NCC differentiation. NCCs are defined as a cell population with multi-potency and also regarded as a fourth germ layer in the developing embryo that contributes to the formation of a wide range of tissues including craniofacial cartilage, heart and pigment cells^43^. Therefore, the compromised differentiation to NCCs upon Hmg20a depletion might also partially account for the observed heart defects. A close examination of cardiac NCCs, a specified subpopulation of NCCs, upon Hmg20a depletion could give a hint to this speculation. Additionally, we have also observed hyperpigmentation in the Hmg20a-depleted tadpoles (see Figure 4A), which might be due to skewed differentiation of NCCs to pigment cells of the skin at the expense of other cell types/tissues. Also, our finding complements a previous report showing the implication of Hmg20a in neuronal development as well as in skeletal muscle differentiation^23, 44^ (see also GO term analysis in Supplemental Figure 6). On the other hand, functional NCCs are marked by their migration ability requiring epithelial to mesenchymal transition (EMT). Here, we demonstrate Hmg20a to be required for EMT in mouse NCCs, which at least partially explains our observed phenotypes. In line with our finding, a previous study reported that HMG20A is required for SNAI1-mediated EMT by replacing HMG20B and its loss induced the reversion of the EMT signalling program^28^.

Regarding the underlying molecular mechanisms during differentiation to NCCs and CMs, HMG20A interacting protein complexes might contribute to the observed defects. BHC/CoREST with KDM1A (LSD1) are well-known factors involved in neural stem cell biology and during neural development^45^. Previous studies suggest the interaction of HMG20A with BHC/CoREST complex plays an important role in the initiation of neuronal differentiation^23, 46, 47^. Recently, the CHD4/NuRD complex was reported to regulate neural differentiation of ESCs^48^. Since neuronal differentiation is associated with NCCs, either BHC/CoREST or NuRD complex might function, together with HMG20A, during NCC differentiation. On the other hand, KDM1A (LSD1) has been shown to play a role in heart development via its interaction with binding partners (e.g. BHC/CoREST) and enzymatic activities^49^. The CHD4/NuRD complex has also recently been demonstrated to directly control cardiac sarcomere formation in the developing heart in mice^50^. Together these studies indicate the potential engagement of HMG20A/CoREST/LSD1 and/or HMG20A/CHD4-NuRD complexes in cardiac development.

Other HMG20A interacting proteins, such as TEADs, also need to be highlighted, as TEAD1 acts as one of the core cardiac transcription factors in heart development^40^. Apart from these interacting partners we identified, it is notable that Yamamoto et al., reported the interaction of HMG20A with Ca2+/S100A6, a protein that contributes to cellular calcium signalling^51^. Very interestingly, we found that *Hmg20a* KO cardiomyocytes are either unable to beat or with much reduced contractility (after a delay) as compared to its WT counterparts. This might be due to the lack of interaction with Ca2+/S100A6, thereby leading to the loss of calcium signalling. In the future, individual knockouts of each HMG20A binding partner in combination with genome-wide profiling of their localisation will be required to further elucidate the role of HMG20A in specific-lineage commitment.

Apart from HMG20A binding partners, we have generated a genome-wide HMG20A chromatin binding map and characterized defined consensus sites (e.g. Jun/Fos), providing a useful resource for follow-up studies. Notably, transcription factors Jun/Fos have been reported to positively bias mESCs to differentiate towards ectoderm^36^, the lineage from which NCCs are derived. This raises the possibility that HMG20A might either compete or indirectly interact with Jun/Fos to regulate gene expression of their downstream targets during differentiation.

In conclusion, our findings implicate HMG20A as part of the H2A.Z/PWWP2A/NuRD- axis that sits at regulatory regions characterized by defined consensus sequences, thereby acting as a key modulator in orchestrating specific transcription programs to ensure proper lineage differentiation.

## Materials & Methods

### Cell culture

HeLa Kyoto (HeLaK) cells were grown in Dulbecco’s modified Eagle’s medium (DMEM, Gibco) supplemented with 10% fetal calf serum (FCS; Gibco) and 1% penicillin/streptomycin (37°C, 5% CO_2_) and were routinely tested for mycoplasma contamination. mESCs were cultivated in 2i + LIF condition as previously described^52^. Specifically, mESCs were cultured on 0.1%-gelatin-coated 6-well plates in N2B27 medium supplemented with 50 μM β-mercaptoethanol, 2 mM L-glutamine, 0.1% Sodium bicarbonate, 0.11% bovine serum albumin fraction V, 1000 units/ml recombinant mouse leukemia inhibitory factor (LIF), 1 µM PD0325901 (MEK inhibitor) and 3 µM CHIR99021 (GSK3 inhibitor). Every two days, cells were passaged at a density of 1.4×10^5^ cells/well. Embryoid body (EB)-mediated differentiation into cardiomyocytes was performed as previously described^38^. Briefly, on differentiation day 0, naïve mESCs (2i+LIF) were adapted to primed state in differentiation media (DMEM, Gibco) supplemented with 10% FCS, 2 mM L-glutamine, 1% nonessential amino acids, 0.1 mM β-mercaptoethanol, 1000 U/mL leukemia inhibitory factor (LIF) for 2 days. Hanging drops containing 1000 cells were prepared in 25 µl of differentiation media (without LIF) supplemented with 50 µg/ml vitamin C. On differentiation day 6 (4 days in suspension), each droplet containing one EB was carefully transferred onto a 0.1%-gelatin-coated 24-well plate. Beating cardiomyocytes were observed starting from differentiation day 7. Transfections were performed using FuGENE® HD Transfection Reagent according to the manufacturer’s instructions (Promega). 24h post-transfection, mESCs were selected by puromycin at a concentration of 0.25 µg/ml for at least 10 days. Finally, following puromycin selection, mCherry-positive mESC colonies were picked for further culture and characterization by RT-qPCR and western blotting, respectively. Sf9 cells were cultured in Sf-900TM II SFM medium (Gibco) and maintained at 27 °C and 90 r.p.m.

### Plasmids

To generate human HMG20A (truncation) fusion proteins, total RNA of HelaK cells was purified using RNeasy-Kit (QIAGEN) and reverse transcripted using Transcriptor First Strand cDNA Synthesis Kit (Roche). cDNA was amplified by Q5-DNA polymerase. (New England Biolabs) and cloned into pIRESneo-GFP^4^ and pFASTBAC1 (Invitrogen) (FLAG was introduced by PCR primer) vectors. To generate hmg20a.L RNA *in situ* hybridization probes, Hmg20a cDNA from *X. laevis* embryos was amplified and cloned into pcDNA3.0. To generate *Hmg20a* KO mESC cell lines CRISPR/Cas9 technology was used. Briefly, sgRNAs were designed to target the start codon (ATG) and the first intron of *Hmg20a* using the online tool (http://crispor.tefor.net), synthesized by Integrated DNA Technology (IDT) and cloned into pX461 vector (Addgene). Donor pUC19-based vectors containing a selectable marker, either mCherry or puromycin resistance gene, and mammalian transcriptional triple terminators bGH+hGH+SV40 (synthesized by GENEWIZ) flanked by homology arms, generated by PCR using Q5-DNA Polymerase from mESC genomic DNA obtained using the QIAamp DNA Mini Kit (QIAGEN), were constructed by HiFi DNA Assembly (New England Biolabs). All relevant sgRNA sequences and primers are listed in Supplemental Table 4.

To perform NuRD binding immunoprecipitations, pcDNA3.1 constructs were prepared that code for the full-length genes for human CHD4 (UniProt ID: Q14839), RBBP4 (UniProt ID: Q09028), MTA1 (UniProt ID: Q13330), GATAD2A (UniProt ID: Q86YP4), MBD2 (UniProt ID: Q9UBB5) ^27^, MBD3 (UniProt ID: O95983), and MTA2 (UniProt ID: O94776), with N-terminal FLAG or HA tags. The exception was full-length HDAC1 (UniProt ID: Q13547) which had the FLAG tag at the C-terminal end^27^. We also used pcDNA3.1 constructs coding for flag-tagged human CHD4 HMG box (residues 1-355), DNA translocase (residues 343-1230), and C-terminal domains (residues 1230-1912).

### siRNA transfections

To deplete HMG20A in HeLaK, 2×10^5^ cells were transfected with 20 pmol of ON-TARGETplus Human HMG20A (10363) siRNA-SMART pool (Dharmacon) using Oligofectamine™ according to the manufacturer’s instructions (Invitrogen). Cells were cultured for 3 days before being harvested for follow-up experiments.

### Antibodies

All used antibodies are listed in Supplemental Table 4.

### Fluorescence Microscopy of HMG20A fusion proteins

1×10^5^ HelaK cells expressing GFP, GFP-HMG20A, GFP-HMG, GFP-CC were seeded onto glass plates and cultured overnight. Next day cells were washed with PBS, fixed for 10 min in 1% Formaldehyde in PBS. After washing fixed cells were permeabilized using 1% bovine serum albumin (BSA) in PBS premixed with 0.1% Triton-X-100 for 30 minutes. Endogenous HMG20A protein was stained by stepwise incubation with primary and then secondary Alexa Fluor–conjugated antibody for 45 min. To visualize DNA cells were treated with 10 µg/ml Hoechst. Coverslips were mounted in Fluoromount-G mounting medium (SouthernBiotech, Birmingham, AL, USA). Images were acquired with an Axio Observer.Z1 inverted microscope (Carl Zeiss, Oberkochen, Germany) with Axiocam 506 mono camera system. Image processing was performed with Zeiss Zen 3.1 (blue edition) software.

### Expression of insect cell Flag-HMG20A and Electromobility Shift Assays (EMSAs)

pFASTBAC1 vectors were transformed into DH10Bac bacteria and viruses were generated according the Bac-to-Bac Baculovirus Expression System protocol (Life Technologies). Extracts from SF9 cells were prepared 3 days after infection by washing the cells twice with ice cold PBS, resuspending and incubation on ice for 10 min in icecold hypotonic buffer (10mM HEPES-KOH pH7.9, 1,5 mM MgCl_2_, 10 mM KCl). After 10 sec vortexing and centrifugation for 10 sec supernatant was aspired. Remaining nuclei were incubated for 20 min in hypertonic buffer C (20 mM HEPES-KOH pH 7.9, 25% Glycerin, 420 mM NaCl, 1,5 mM MgCl_2,_ 0.2 mM EDTA). After centrifugation for 10 min supernatant was either frozen at -20°C or directly used for EMSAs. EMSAs were performed as initially described ^53^ with the exception that 400 ng salmon sperm DNA per reaction was used as unspecific competitor. As EMSA- probe was used the Cy5 end-labelled double-stranded oligo: CAGGGCTAGTGGATCCCNNNNNNNNNNNNNNTGATTCTGTGGATAACCGTATTACCGCCTTTGAGTGAGCTGATACCGCTCGCGGGCTGCAGGAATTCGA

### Preparation of nuclear extracts or S1 mononucleosomes and immunoprecipitation

For each IP, nuclear extracts containing soluble mononucleosomes upon micrococcal nuclease digestion from 2 x 10^7^ GFP-HMG20A expression HelaK cells were prepared and purified as previously described^14^. Purified protein complexes were either subjected to label free quantitative mass spectrometry or Immunoblot analyses. Information about used antibodies are listed in Supplemental Table 4.

### Label-free quantitative Mass-spectrometry (lf-qMS)

The sample preparation of HK cell-derived mononucleosomes and following label-free quantitative mass-spectrometry were carried out as previously described^14^ with the following difference: on-bead tryptic digest was performed using Trypsin Gold, Mass Spectrometry Grade (Promega Corporation, Madison, WI, USA).

#### LC-MS/MS analysis

Peptides were analysed by reversed-phase liquid chromatography on an EASY-nLC 1000 or 1200 system (Thermo Fisher Scientific, Odense, Denmark) coupled to a Q Exactive plus or HF mass spectrometer (Thermo Fisher Scientific). HPLC columns of 50cm length and an inner diameter of 75µm were in-house packed with ReproSil-Pur 120 C18-AQ 1.9µm particles (Dr. Maisch GmbH, Germany). Peptide mixtures were separated using linear gradients of 120 or 140 minutes (total run time + washout) and a two-buffer system: buffer A++ (0.1% formic acid) and buffer B++ (0.1% formic acid in 80% acetonitrile). The mass spectrometer was operated in a data-dependent top 10 or top 15 mode. Peptides were fragmented by higher energy collisional dissociation (HCD) with a normalized collision energy of 27.

#### MS Data analysis

MS raw data were processed using the MaxQuant software version 1.4.3.13^54^. Fragmentation spectra were searched against a human sequence database obtained from Uniprot in May 2013 and a file containing frequently observed contaminants such as human keratins. Cysteine carbamidomethylation was set as a fixed modification; N-terminal acetylation and methionine oxidation were set as variable modifications. Trypsin was chosen as specific enzyme, with 2 maximum missed cleavages allowed. Peptide and protein identifications were filtered at a 1% FDR. Label-free quantification was performed using the MaxLFQ algorithm^54^ integrated into MaxQuant. The match between runs option was enabled with a matching time window of 0.5 min and an alignment time window of 20 min. All other parameters were left at standard settings. MaxQuant output tables were analysed in Perseus^55^ version 1.5.8.6 as follows: After deleting proteins only identified with modified peptides, hits to the reverse database, contaminants and proteins with one or less razor and unique peptides, label-free intensities were log2 transformed. Next, proteins were required to have 3 valid values in at least one triplicate, then remaining missing values in the data matrix were imputed with values representing a normal distribution around the detection limit of the mass spectrometer. Now a two-sample t-test was performed to identify proteins enriched in the HMG20A pull-downs compared input control. Only those proteins were kept for further analysis. The S0 and FDR parameters were set to 0.5 and 0.05, respectively (Supplemental Table 1).

## NuRD binding studies

### HEK293 cell protein expression

Experiments were performed as described^27^. In brief, suspension-adapted HEK Expi293F^™^ cells (Thermo Fisher Scientific, Waltham, MA, USA, Cat no. #A14527) were co-transfected with combinations of plasmids. Cells were incubated for 65 h at 37 °C with 5% CO_2_ and shaking at 130 rpm, then centrifuged (300 *g*, 5 min) and stored at −80 °C.

### Lysate preparation and immunoprecipitation of GFP-HMG20A and NuRD members

Cell extracts were prepared based on a modified version of a previously described protocol^27^. In brief, cell pellets were lysed as described previously and the lysate was then clarified via centrifugation (20,000 *g*, 20 min, 4 °C); the cleared supernatant was used for GFP-affinity pulldowns as described below.

To prepare GFP-beads, streptavidin beads (Thermo Fisher Scientific, Waltham, MA, USA) were first loaded with 6xHis-SUMO-strep-tag-GFP-nanobody protein expressed and purified from *E. coli* BL21 cells. Immobilized GFP nanobody on beads captures soluble GFP-HMG20A and other proteins interacting with it. Cleared supernatant samples were mixed with 20 μL of streptavidin beads pre-loaded with 3 μg of GFP-nanobodies and incubated with rotation for 2 h at 4 °C. Post-incubation, the beads were first washed with 3 × 1 mL wash buffer (50 mM HEPES/NaOH, 500 mM NaCl, 0.5% (v/v) IGEPAL® CA630, 3 mM ATP, 0.2 mM DTT, pH 7.5) and then 2 x 1 mL wash buffer (50 mM HEPES/NaOH, 150 mm NaCl, 0.5% (v/v) IGEPAL® CA630, 0.2 mM DTT, pH 7.5). Bound proteins were eluted by 3 × 20 μL treatment with elution buffer (20 mM HEPES/NaOH, 150 mM NaCl, 100 mM biotin, 0.2 mM DTT, pH 8). Eluted fractions were pooled to be analysed by western blot. Gels were blotted onto PVDF membrane, blocked for 1 h in PBS-T containing 10% (w/v) skim milk and incubated overnight at 4 °C with HRP-conjugated antibodies diluted in PBS-T containing 2% (w/v) skim milk powder. After washes, membranes were imaged using ECL Western Blotting Detection Reagent (GE Healthcare).

### ChIP-seq

ChIP-seq and ChIP-qPCR was performed as previously described^56^ with the difference that 1 x 10^7^ cells were crosslinked in 2 ml culturing medium with 1% formaldehyde. Information about used qPCR primers are listed in Supplemental Table 4.

#### Analysis of ChIP-seq and CUT&RUN data

Image analysis and base calling were performed using the Illumina pipeline v 1.8 (Illumina Inc.)^57^.

Public data sets used in this study:

H3K4me3, PWWP2A, H2A.Z.1 and H2A.Z.2 data from HeLaK cells were used as previously deposited at GEO (GSE78009). ChIP-seq data for additional histone modifications was downloaded from the ENCODE portal at UCSC (http://hgdownload.soe.ucsc.edu/goldenPath/hg19/encodeDCC/wgEncodeBroadHist <u>one/)</u>.

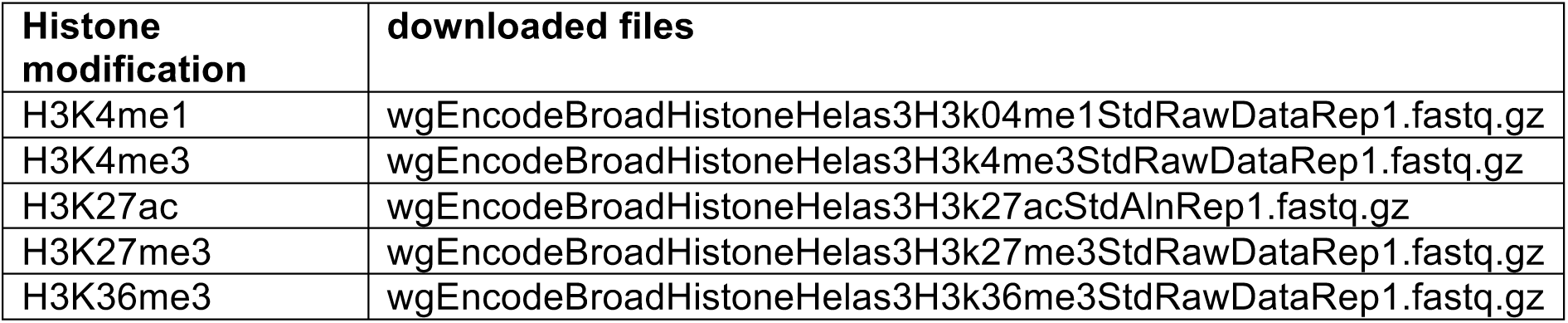

*Plotting and statistics*^58, 59^

Manipulation of sequencing reads was done using Rsamtools^60^ and genomic intervals were represented as GenomicRanges objects^61^. The analysis of the association between peak intervals and known genomic annotation feature were done using the ChIPseeker package^62^ with default setting using the UCSC hg19 gene definitions (BioConductor package TxDb.Hsapiens.UCSC.hg19.knownGene). As statistical tests, we performed Wilcoxon rank sum tests. The code underlying our analysis is available upon request.

## RT-qPCR and RNA-seq

RNA isolation, RT-qPCR and RNA sequencing was performed as previously described^56^. With the exception of RNA sequencing of mESCs, which was performed at Novogene (UK). Information about used qPCR primers are listed in Supplemental Table 4.

### RNA-seq analysis

Trimming was performed identical to the CUT&RUN data. Alignment of the trimmed FASTQ files against the mm9 genome (or hg19 for HeLa data) was performed using hisat2 v.2.2.1 with “--min-intronlen 30--max-intronlen 3000” parameters. The following analysis steps were performed within R v.4.1.2^63^ using a modified version of R/BioConductor package systemPipeR^64^ for various steps. Based on the BAM files and the mouse mm9 GTF (or hg19 GTF for HeLaK data) read counts per gene for each sample were calculated using the summarizedOverlaps function of the GenomicAlignments^61^ R package. The resulting read counts were normalized using DESeq2 v.1.28.1^65^. DESeq2 was used for the identification of differentially expressed genes (log2FC > 2 or log2FC < -2 for mESCs and log2FC > 0.8 or log2FC < -0.8 for HeLaK and adjusted p-value < 0.05) for the displayed contrasts, unless otherwise indicated. The PCA was calculated using DESeq2 and plotted using ggplot2^66^. The z-scaled heatmap was clustered according to the Euclidian distance using the “ward.D2” method. Line plots for the gene expression at different days were min-max normalized based on all expression values for each gene. Snapshots based on the coverage tracks were generated using the Gviz package^59^. Gene ontology analysis for the genes of different clusters was performed using Metascape^67^ web interface (www.metascape.org) and plotted using ggplot2.

### Primers

See Supplemental Table 4.

### Xenopus laevis experiments

*Xenopus laevis* staging, microinjection, lacZ staining and whole mount *in situ* hybridization were performed as previously described^68^. For microinjections, capped sense mRNA *(lacZ, mbGFP*^69^ and *mbRFP*^70^) and the following Morpholino Oligonucleotides (MO) were used: standard control morpholino (co MO, 5 ′-CCTCTTACCTCAGTTACAATTTATA-3′, Gene Tools, LLC) and hmg20a translation blocking MO (hmg20a MO, 5′- TGCAGAGGCTGTGCTTTCCATCTAG-3′, Gene Tools, LLC). The spatial *hmg20a* expression pattern was characterized using albino embryos; sense controls were analysed for all documented stages. Histological sections were obtained as previously described^71^. For phenotypical characterization of craniofacial and heart structures, 80 pg *lacZ* RNA, 50 pg *mbGFP* RNA or 80 pg *mbRFP* RNA were co-injected as lineage tracers to mark the injected side. Cartilage development was analysed by whole-mount immunofluorescence staining as previously described^72^. Cartilage phenotypes were quantified by measuring the area of the ceratohyal cartilage using the polygon function of ImageJ. The ratio of the relative surface area of the Morpholino-injected side and the control side was calculated and plotted in a boxplot diagram. For phenotypical and immunofluorescence documentation, a Nikon stereo microscope (SMZ18) with a DS-Fi3 Nikon Camera and the NIS-Elements imaging software was used.

## CUT&RUN-seq

### Preparation of samples

CUT&RUN was performed on 5×10^5^ mESCs at primed stage of cardiomyocyte differentiation protocol (Figure 5D) applying the CUTANA® CUT&RUN Kit (version 2) according to manufacturer instructions. Information about used antibodies are listed in Supplemental Table 4. CUT&RUN sequencing libraries were generated using NEBNext® Ultra™ II DNA Library Prep Kit for Illumina® (New England Biolabs) according to manufacturer instruction. Sequencing, was performed at Novogene (UK). *Bioinformatic analysis*

Paired end raw FASTQ files were quality and adaptor trimmed using trimGalore v.1.18^73^. Trimmed FASTQ files were aligned against the mouse mm9 reference genome (Illumina’s iGenomes) using hisat2 v.2.2.1^74^ with the “--no-spliced-alignment” parameter and stored as Binary Alignment Map (BAM) files. PCR duplicated reads were removed from BAM files using Picard tools v.2.21.9 (http://picard.sourceforge.net). The resulting BAM files were used to generate individual coverage tracks (bigWig) for each sample using deepTools bamCoverage function^75^. MACS2 v.2.2.7.1^76^ with IGG from wild type or knock out as an input was used for the peak calling on the two wild type and two Hmg20a knock out samples. Only peaks from the wild type samples that were not identified in one of the Hmg20a KO samples were used as the real Hmg20a binding sites. Additionally, those sites were filtered for known mouse mm9 blacklisted regions^77^. Based on those 2545 bona fide Hmg20a sites and the individual coverage tracks for each sample deepTools computeMatrix and plotHeatmap commands were used to generate the binding heatmap. ChIPseeker^62^ with the USCSC’s mm9 Gene transfer format (GTF) files was used to identify the genomic features that are associated with Hmg20a binding sites. MEME-Suite was used for the motif discovery analysis of the Hmg20a binding sites.

## Acknowledgement

We thank Sonja Sahner and Lena Vogel for technical help and all current and past Hake group members for theoretical and practical advice. This work was supported by the Deutsche Forschungsgemeinschaft (DFG) through the Collaborative Research Center TRR 81 (projects A12, A15, and Z1) to TB, SBH and MB, respectively and the Cardio Pulmonary Institute (CPI) to SBH. SG was funded by the DFG Research Training Group GRK 2213, Membrane Plasticity in Tissue Development and Remodeling. The work was also supported by a grant from the National Health and Medical Research Council of Australia to JPM (APP1126357).

## Author contributions

AH, JL and SBH conceived this study. AH cloned all constructs, performed IP, ChIP-seq, CUT&RUN and RNA preparations with the help of FD, JL, IS and LVZ. ND executed microscopy pictures and FACS analyses with the help of AH. JLe generated recombinant FLAG-HMG20A proteins in Sf9 cells and performed EMSAs. SG performed all *Xenopus* experiments with support of LS, SGe and AB. AR executed lf- qMS identification with support of MM. Next generation sequencing of ChIP samples was applied by AN with support of TS. All bioinformatics analyses were performed by TF and MB, with support of TB. NuRD member binding studies were carried out by HMS and JPM. JL and AH performed all mESC differentiation experiments. SBH, AB, MB, JL and AH wrote the manuscript with support from all other co-authors.

