## Supplemental Material for "The H2A.Z.1/PWWP2A/NuRD-associated protein HMG20A controls early head and heart developmental transcription programs"

**to**

Supplementary Figure 1

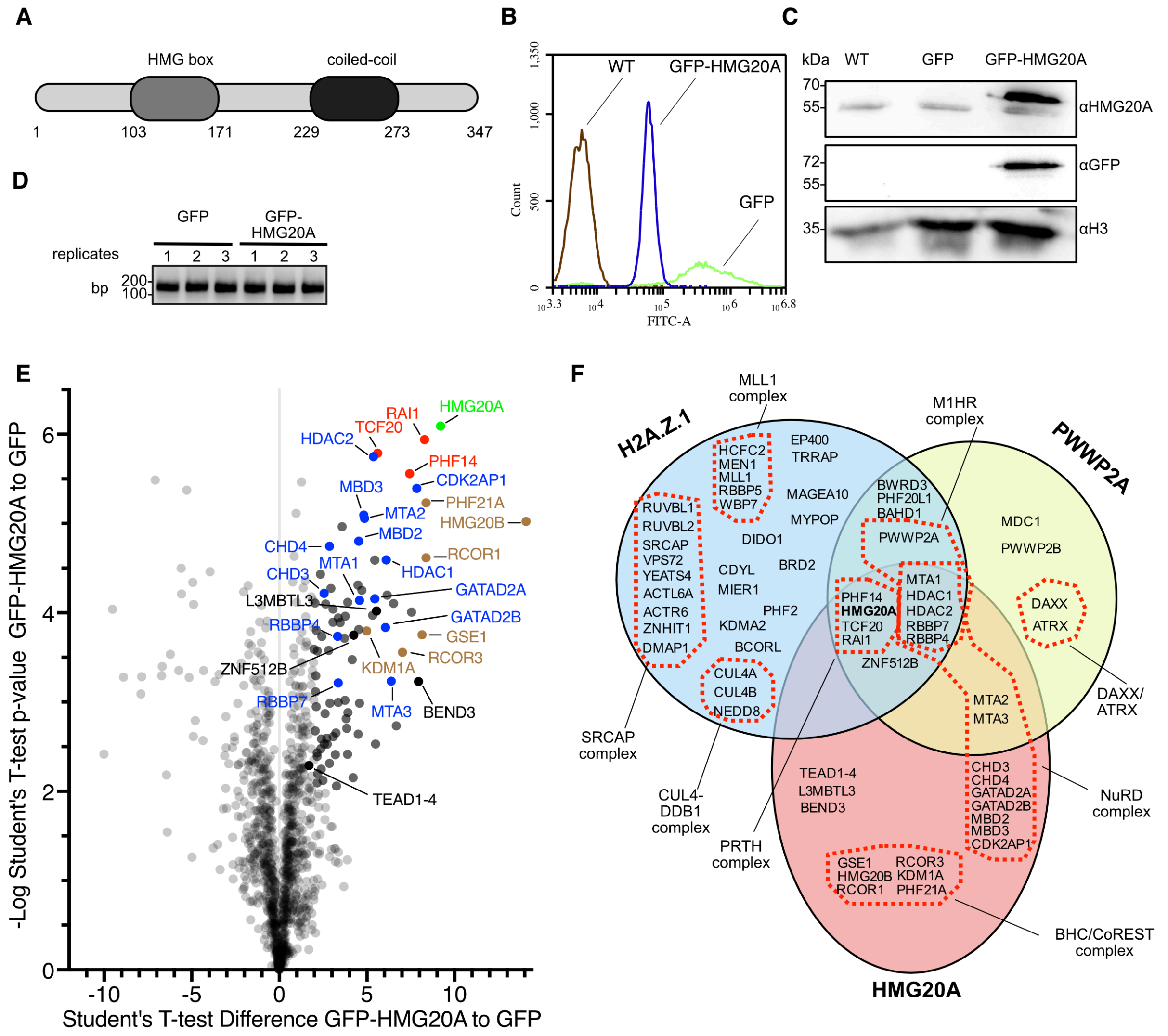


**Supplemental Figure 1: HMG20A binds chromatin modifying complexes.**

**(A)** Schematic depiction of human HMG20A protein with its N-terminal HMG box and C-terminal coiled-coil (CC) domain.

**(B)** Flow cytometry analysis of HeLaK cells (WT, brown) stably expressing GFP (green) or GFP-HMG20A (blue).

**(C)** Immunoblot of cell extracts from HeLaK cells (WT) stably expressing GFP or GFP-HMG20A with anti-GFP and anti-HMG20A antibodies. Anti-H3 serves as loading control.

**(D)** Agarose gel of purified DNA fragments after MNase digestion using HeLaK cells stably GFP and GFP-HMG20A.

**(E)** Volcano plot of second replicate of label-free interaction partners of GFP-HMG20A-associated mononucleosomes. Significantly enriched proteins over GFP-associated mononucleosomes are shown in upper right part. T-test differences were obtained by two-sample t-test. HMG20A is highlighted in bright green, PRTH members in red, BHC/CoREST members in brown, NuRD members in blue, other proteins in black and background binding proteins in grey. See also Figure 1C for Volcano plot of first biological replicate and Supplemental Table 1 for detailed list of HMG20A binders.

**(F)** Schematic depiction of overlapping H2A.Z.1 [1,2], PWWP2A [3] and HMG20A interactomes.

#### Supplementary Figure 2

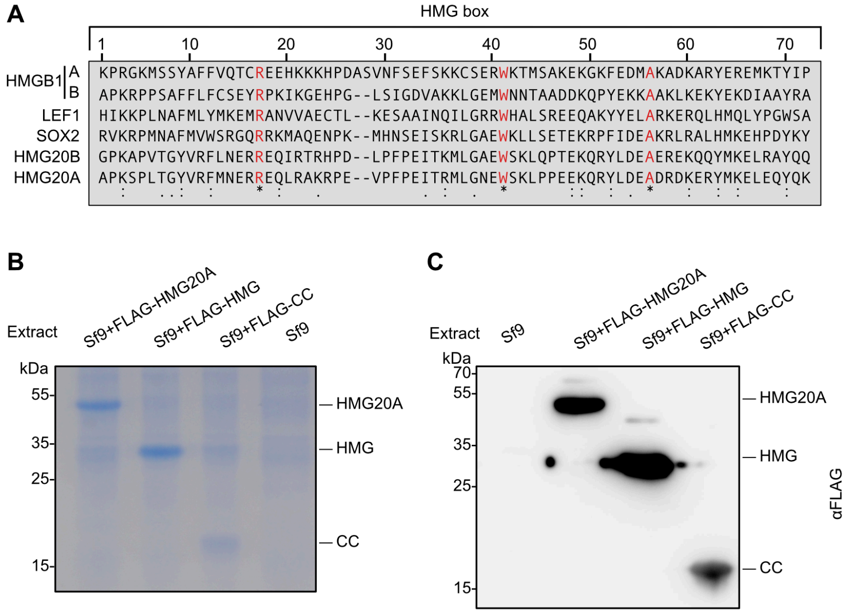


**Supplemental Figure 2: Expression and purification of HMG20A deletion proteins.**

**(A)** Alignment of HMG box amino acid sequences from diverse human HMG-box containing proteins. Alignment was performed using Clustal Omega.

**(B, C)** Coomassie-stained SDS-PAGE gel (**B)** or anti-FLAG immunoblot **(C)** of extracts from Sf9 cells expressing FLAG-HMG20A, -HMG or -CC proteins.

Supplementary Figure 3

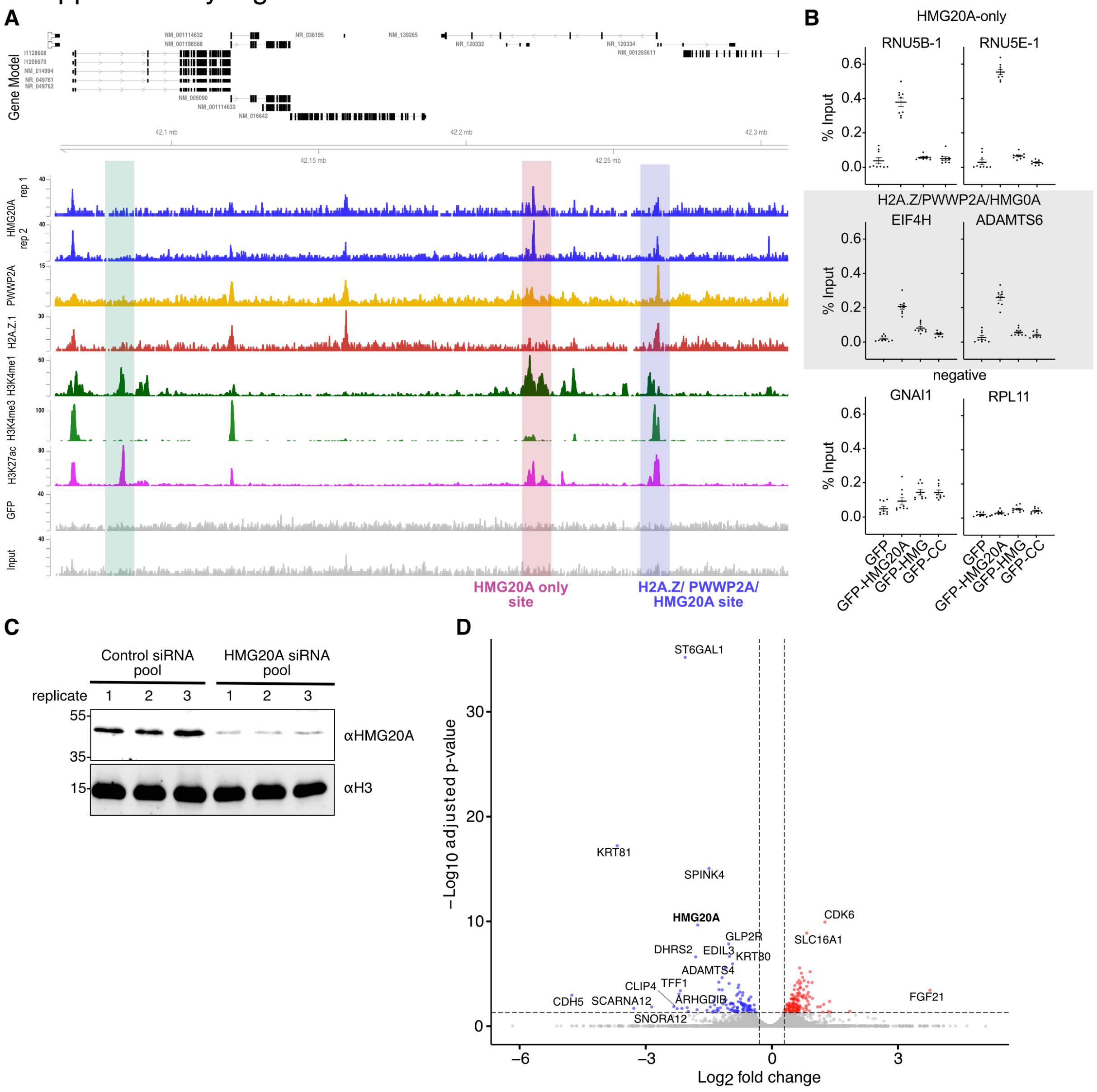


**Supplemental Figure 3: HMG20A binds regulatory genomic regions but has no major transcriptional impact in HeLaK cells**

**(A)** Genome browser snapshot of a representative region in chromosome 15 displaying input (grey), GFP control (grey), H3K27ac, (purple), H3K4me3 (light green), H3K4me1 (dark green), GFP-H2A.Z.1 (red), GFP-PWWP2A (orange) and two replicates of GFP-HMG20A (blue) ChIP-seq signals. Blue bar depicts H2A.Z.1/PWWP2A/HMG20A-positive site, red bar depicts HMG20A-only site and green bar depicts a H2A.Z.1/PWWP2A/HMG20A-negative control site.

**(B)** Validation of ChIP-seq data by ChIP-qPCR at selected loci. Shown is percent input of three replicates of GFP, GFP-HMG20A, -HMG or -CC ChIP-qPCR of HMG20A-only sites (RNUB-1 and RNUE-1downstream; see red bar in A as example),

H2A.Z.1/PWWP2A/HMG20A-positive sites (EIF4H promoter and ADAMTS3 gene body; see blue bar in A as example) and an H2A.Z.1/PWWP2A/HMG20A-negative site (RPL11 gene body; see green bar in A as example) as negative control. Error bars indicate SEM (n=9).

**(C)** Immunoblot of HMG20A upon siRNA-mediated depletion in HelaK cells.

**(D)** Volcano plot of significantly deregulated (log2 fold change <-1, p<0.05) mRNAs from two independent siRNA-mediated HMG20A depletion experiments analysed by RNA-seq. Red: upregulated transcripts, blue: downregulated transcripts.

### Supplemental figure 4

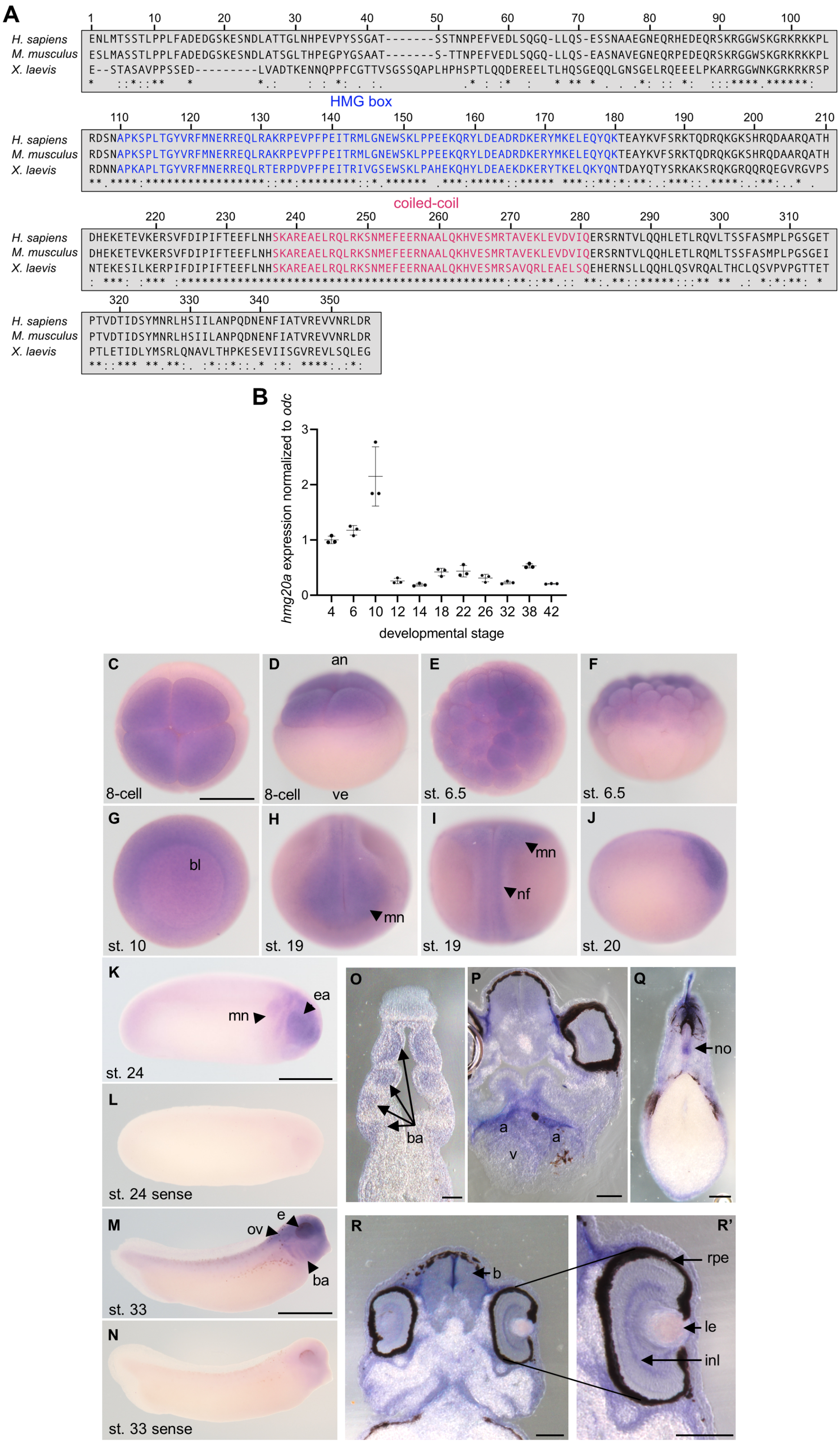


**Supplemental Figure 4: *hmg20a* expression in *X. laevis*.**

**(A)** Alignment of human (*H. sapiens*), mouse (*M. musculus*) and frog (*X. laevis*) Hmg20a protein sequences. HMG box is shown in blue, coiled-coil region in red. Alignment was performed using Clustal Omega.

**(B)** Temporal expression pattern of *Xenopus hmg20a:* RT-qPCR of *hmg20a* mRNA expression covering *X. laevis* developmental stages 4 (8-cell stage) to 42 normalized to *odc* expression. Error bars indicate s.e.m. of three technical replicates.

**(C-R')** Spatial expression pattern of *hmg20A* determined by whole mount *in situ* hybridization. *hmg20A* mRNA is detected at early stages of *Xenopus laevis* development. **(C)** 8-cell stage embryo, anterior view. **(D)** 8-cell stage embryo, dorso-lateral view, animal and vegetal pole are indicated. **(E)** Embryo at blastula stage 6.5. anterior view. **(F)** Same embryo as in **E**, dorsal view. **(G)** Embryo at gastrula stage 10. **(H)** Embryo at neurula stage 19, anterior view. **(I)** Same embryo as in **H**, dorsal view. **(J)** Embryo at stage 20, lateral view. **(K)** Embryo at stage 24, lateral view. **(L)** Sense control, embryo at stage 24. **(M)** Embryo at stage 33, lateral view. **(N)** Sense control, embryo at stage 33. Scale bar in **C-N** is 1mm. **(O)** Transverse section through the branchial arch region of a stage 31 embryo, *hmg20A* expression in the branchial arches is indicated by arrows. **(P-R')** Transverse sections of a stage 42 embryo. **(P)** *hmg20A* is partially expressed in the heart region. **(Q)** *hmg20A* expression within the notochord (no). **(R, R')** *hmg20A* is partially expressed in the brain and eye. Scale bar in **O-R'** is 100 µm. abbreviations: a, atrium, an, animal; b, brain; ba, branchial arches; bl, blastoporus; ea, eye anlage; e, eye; inl, inner nuclear layer; le, lens; mn, migratory neural crest; nf, neural fold; no, notochord; ov, otic vesicle; rpe, retinal pigment epithelium v, ventricle; ve, vegetal.

Supplementary Figure 5

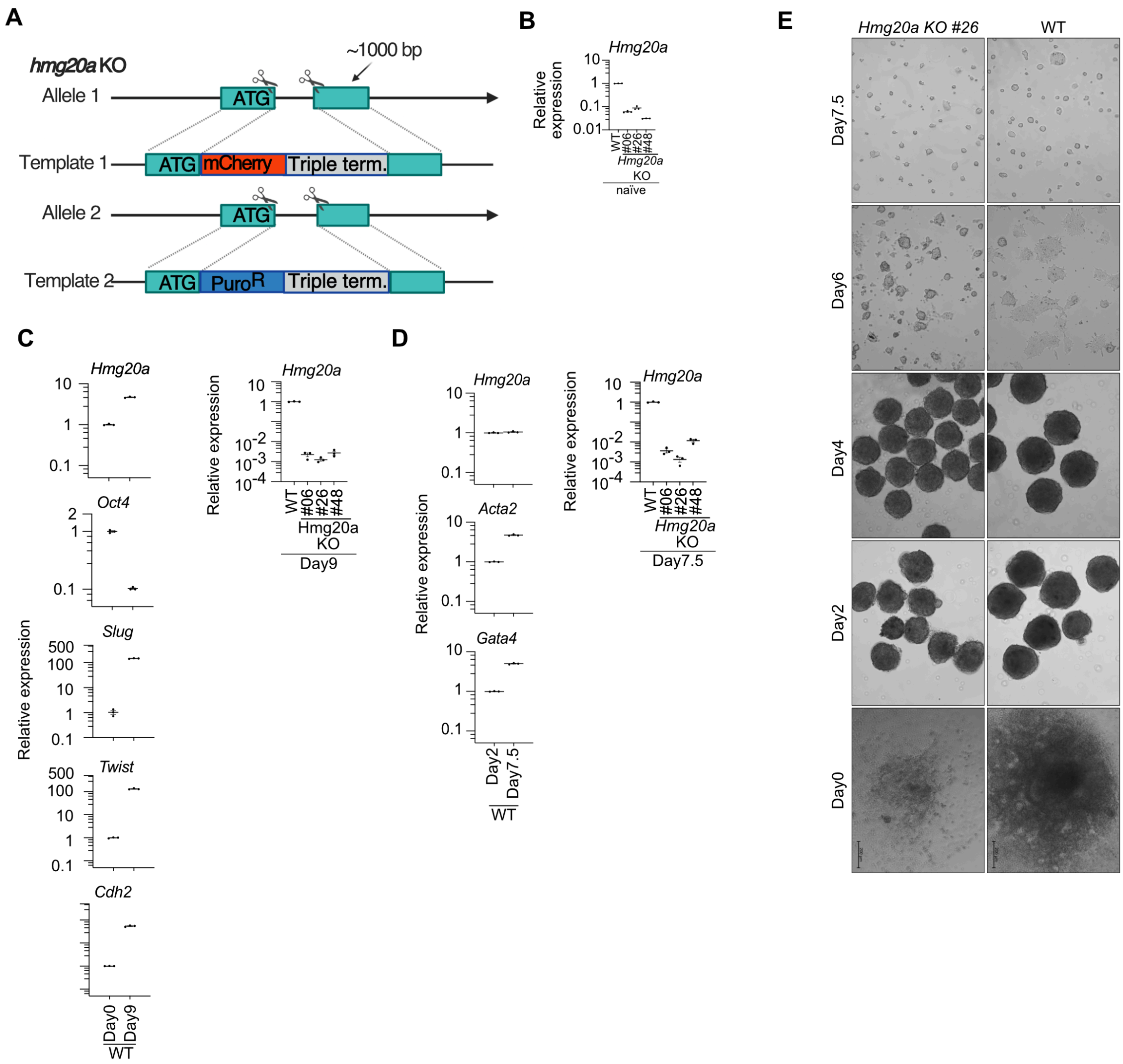


**Supplemental Figure 5: Loss of Hmg20a impairs mESC differentiation**

**(A)** Schematic depiction of *hmg20A* KO generation in mESCs by introducing mCherry_triple terminator sites and a puromycin resistance_triple terminator site into both *hmg20a* alleles directly after the start codon by CRISPR/Cas9-based approach.

**(B)** Validation of three *hmg20A* KO clones by RT-qPCR. Shown is *hmg20* expression normalized to *Hprt*. Error bars indicate SEM of three technical replicates.

**(C)** Left: RT-qPCR of EMT marker genes *Slug, Twist Cdh2,* pluripotency marker *Oct4* and *hmg20a* in WT cells at Day0 and Day9 of neural crest differentiation protocol. Right: RT-qPCR of *hmg20a* in WT and 3 individual *hmg20a* KO clones at Day9 of neural crest differentiation protocol. Expression was normalized to *Hprt, 16S RNA and Gapdh* expression.

**(D)** Top: RT-qPCR of *hmg20a* (left) and cardiomyocyte marker genes *Acta2* (middle) *and Gata4* (right) in WT cells at Day2 and Day7.5 of neural crest differentiation protocol. Bottom: RT-qPCR of *hmg20a* in WT and 3 individual *hmg20a* KO clones at Day7.5 of cardiomyocyte differentiation protocol. Expression was normalized to *Hprt, 16S RNA and Gapdh* expression.

**(E)** Phase-contrast microscopy images of WT and *hmg20a* KO mESCs during cardiomyocyte differentiation. Scale bar: 200 µm

Supplementary Figure 6

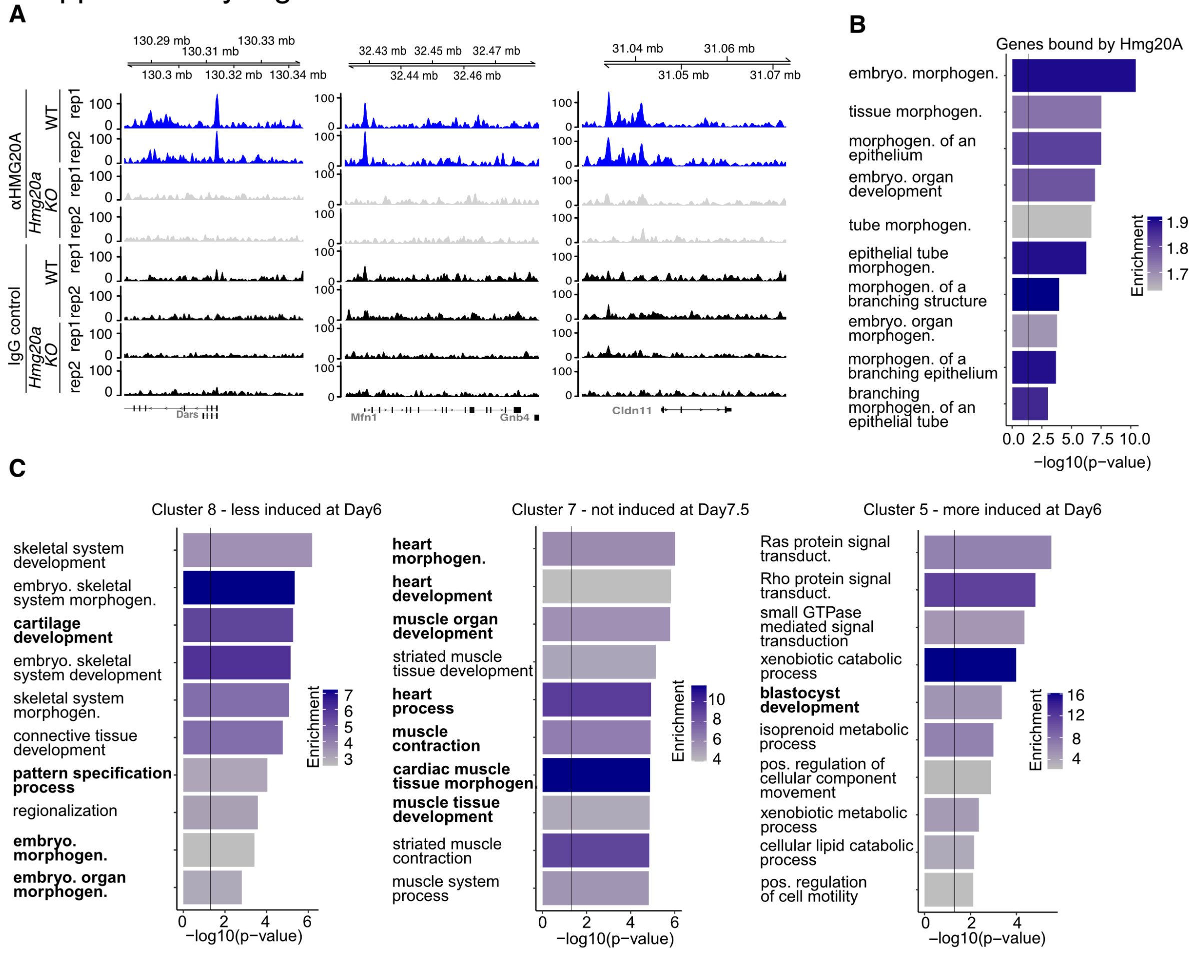


**Supplemental Figure 6: Hmg20a regulates cardiomyocyte transcription programs**

**(A)** Genome browser snapshot of representative Hmg20a binding regions. GO term analysis of genes bound by Hmg20a **(B)** and of genes in critical gene clusters identified in Figure 6F **(C)**.
